# Pre-mRNA splicing order is predetermined and maintains splicing fidelity across multi-intronic transcripts

**DOI:** 10.1101/2022.08.12.503515

**Authors:** Karine Choquet, Autum Koenigs, Sarah-Luisa Dülk, Brendan M. Smalec, Silvi Rouskin, L. Stirling Churchman

## Abstract

Combinatorially, intron excision within a given nascent transcript could proceed down any of thousands of paths, each of which would expose different dynamic landscapes of cis-elements and contribute to alternative splicing. In this study, we found that post-transcriptional multi-intron splicing order in human cells is largely predetermined, with most genes spliced in one or a few predominant orders. Strikingly, these orders were conserved across cell types and stages of motor neuron differentiation. Introns flanking alternatively spliced exons were frequently excised last, after their neighboring introns. Perturbations to the spliceosomal U2 snRNA altered the preferred splicing order of many genes, and these alterations were associated with the retention of other introns in the same transcript. In one gene, early removal of specific introns was sufficient to induce delayed excision of three proximal introns, and this delay was caused by two distinct cis-regulatory mechanisms. Together, our results demonstrate that multi-intron splicing order in human cells is predetermined, is influenced by a component of the spliceosome, and ensures splicing fidelity across long pre-mRNAs.

## Introduction

Pre-mRNA splicing is an essential regulatory step of gene expression in which introns are removed and exons are ligated to generate mature mRNAs. Human genes contain many introns, and more than 95% of these genes undergo alternative splicing (AS)^1,2^. Accordingly, pre-mRNA splicing requires intricate mechanisms to coordinate the excision of multiple introns within each nascent transcript. Furthermore, the timing of splicing catalysis varies across introns. Sequencing studies have shown that up to 70% of introns are removed co-transcriptionally^3,4^, whereas the other 30% are excised post-transcriptionally while the nascent transcripts remain associated with chromatin after 3’-end cleavage and polyadenylation^3,5,6^. Although mRNA isoforms within the mature transcriptome have been explored in many cell types and tissues^7,8^, it remains unclear how splicing and AS decisions occur across multiple introns of a single transcript to produce these pools of mature isoforms.

The order of intron removal is emerging as a crucial regulatory component of splicing. Because many splicing enhancers and silencers are located within introns^9–12^, the order of intron excision determines how long these elements persist within a transcript, where they can influence AS outcomes in cis. How splicing order proceeds across multiple introns within a single transcript remains largely unexplored, and the problem is immense: for the average human gene with eight introns, more than 40,000 possible paths can lead from an unspliced pre-mRNA to a fully spliced transcript. Due to technical limitations of analyzing splicing patterns across many introns, global analyses of intron removal order have thus far been limited to pairs of consecutive introns. These studies have shown that removal occurs largely in a defined order, i.e., one intron of a pair is usually excised before the other^13,14^. Multi-intron splicing order across more than two introns has been analyzed for a few individual genes by RT-PCR or targeted next-generation sequencing^15–18^, but because these experiments require complex primer design, they are limited to analyses of a handful of genes. Long-read transcriptome sequencing has emerged as a promising method for investigating the order of RNA processing events within single RNA molecules^14,19,20^. However, the low throughput of long-read technologies limited results to aggregate analyses of all introns across all sequenced transcripts. These initial studies have demonstrated that neighboring introns are more likely to have the same excision status than more distant introns, suggesting that removal of proximal introns is coordinated^14^, but it is unclear whether this coordination extends to larger groups of introns within a single transcript and how it impacts AS outcomes.

Splicing is performed by the spliceosome, a complex consisting of five small nuclear ribonucleoproteins (snRNPs) that each contain one small nuclear RNA (snRNA) and several proteins^21^. Early in spliceosomal assembly, the 5’ splice site (SS) and branch point sequence (BPS) in the pre-mRNA interact with the U1 and U2 snRNAs via sequence complementarity. The relative levels of U1 and U2 snRNAs influence splicing fidelity, as demonstrated by the observation that moderate depletion of U1 and U2 snRNAs causes AS changes that are enriched for exon skipping events^22^. In mice, a mutation in only one of the five genes encoding U2 snRNA is sufficient to cause significant exon skipping and induces neurodegeneration^23^. We previously found that introns that are excised later within pairs are enriched for binding of U2 snRNP components^14^. Moreover, the U2 snRNP is important for synergistic spliceosome assembly across proximal introns *in vitro*^24^, suggesting a role for the U2 snRNP in regulating splicing order and coordination.

In this study, we used direct RNA nanopore sequencing to study multi-intron splicing order during post-transcriptional splicing. We found that post-transcriptional splicing is widespread in human cells and frequently occurs in several introns per gene. In addition, we observed that post-transcriptional splicing across three or four proximal introns follows a defined order that is conserved across cell types and stages of human motor neuron differentiation, indicating that splicing order is generally predetermined and not subject to substantial variation. Moreover, depletion of U2 snRNA or targeted perturbation of selected introns using antisense oligonucleotides led to changes in splicing order that were accompanied by splicing defects throughout the resulting splice isoform. For example, out-of-order removal of one intron in the gene *IFRD2* was sufficient to increase the retention of several proximal introns through two distinct cis-regulatory mechanisms involving a downstream SS or changes in secondary structure. Together, our results demonstrate that multi-intron splicing order in human cells is predetermined, regulated by a core component of the spliceosome and pre-mRNA *cis*-elements, and crucial for splicing fidelity.

## Results

### Analysis of post-transcriptional splicing by direct RNA sequencing

To investigate post-transcriptional splicing across entire transcripts, we used nanopore direct RNA sequencing to analyze chromatin-associated poly(A)-selected RNA from human K562 cells (Fig. 1A). We focused on post-transcriptional splicing because we previously showed that co-transcriptional splicing occurs several kilobases away from transcription^14^, making it challenging to analyze with the current nanopore sequencing read lengths, as most reads corresponding to elongating transcripts are completely unspliced (Extended Data Fig. 1A). Moreover, introns neighboring alternative exons are predominantly excised post-transcriptionally^3^ and are more likely to be removed second within intron pairs^14^, suggesting that studying post-transcriptional splicing will shed light on AS.

**Figure 1.**
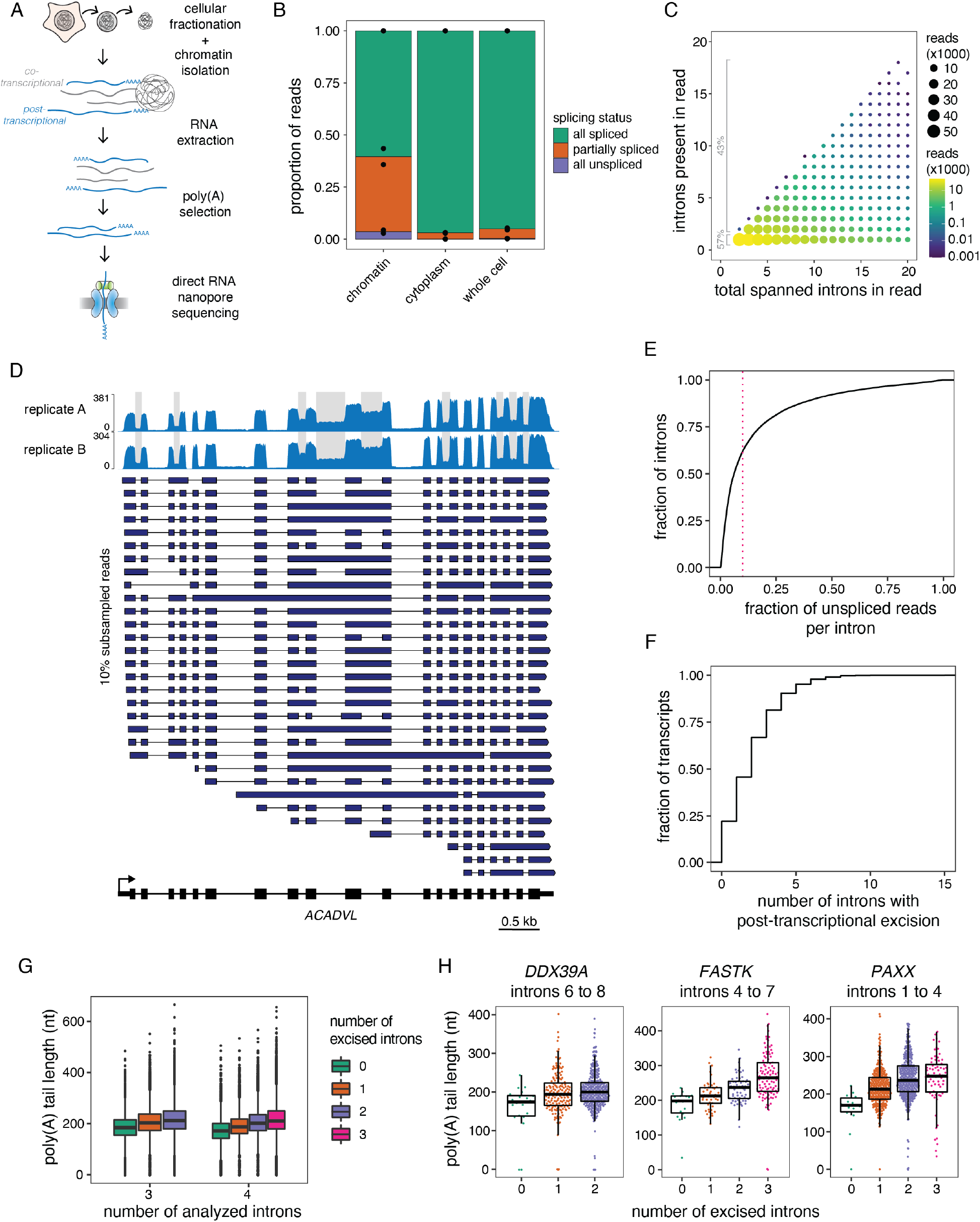
Widespread post-transcriptional splicing in human cells. A) Schematic depicting the experimental design. Cellular fractionation was performed on human K562 cells to isolate the chromatin, followed by purification of chromatin-associated RNA. Polyadenylated RNAs were enriched using oligo(dT) beads and underwent direct RNA nanopore sequencing. B) Proportion of reads spanning at least two introns that are fully spliced, partially spliced (intermediate isoforms), or fully unspliced in direct RNA sequencing samples from different cellular compartments. Individual dots represent two biological replicates. C) Post-transcriptionally excised introns in direct RNA sequencing of poly(A)-selected chromatin-associated RNA. The number of present introns in each read is represented as a function of the total number of introns spanned by the read (excised or not). The size and color of the dots indicate the number of reads in each category. The percent of reads with one intron present and with more than one intron present is shown in grey font. D) Representative example gene displaying post-transcriptional splicing. Introns with fraction unspliced reads > 0.1 are shaded in grey. The overall coverage track is shown for two biological replicates at the top and 10% randomly sampled reads from replicate B are shown below as dark blue arrows. The gene structure is shown at the bottom, with rectangles representing exons and lines indicating introns. The arrow indicates the transcription start site. E) Cumulative distribution function (CDF) showing the fraction of unspliced reads per intron. The red dotted line indicates 10% or more unspliced reads and represents the cutoff used to define post-transcriptional splicing. F) CDF showing the number of post-transcriptionally excised introns per gene. For E) and F), only reads spanning at least two introns were included. G) Distribution of poly(A) tail lengths in chromatin-associated RNA from K562 cells. All reads are classified into groups of 3 or 4 introns based on the number of proximal post-transcriptionally excised introns that the read covers. Each intron group must have at least 10 intermediate isoform reads for each number of excised introns in the read. H) Same as G, but for example individual intron groups. Each dot represents one read. In G) and H), boxplots elements are shown as follows: center line, median; box limits, upper and lower quartiles; whiskers, 1.5x interquartile range; In G), points represent outliers.

Using our dataset of 2.65 million aligned reads (Supplemental Table 1), we classified the splicing status of each read spanning at least two introns (1.59 million reads) as being fully spliced, fully unspliced, or partially spliced (hereafter, intermediate isoform reads). More than 35% of multi-intron reads exhibited partial splicing, likely reflecting transcripts that were undergoing RNA processing, compared to less than 5% for cytoplasmic and total RNA (Fig. 1B). We observed that 57% of intermediate isoform reads in chromatin RNA contained only one remaining intron, whereas 43% had two or more remaining introns (Fig. 1C). The majority of introns that were not excised were middle introns (i.e., not the first or last introns in the transcript) (Extended Data Fig. 1B). Our analysis captured 66% of genes expressed in K562 cells (RPKM > 1). We found that 38% of introns were excised post-transcriptionally, defined as being present in at least 10% of reads (Fig. 1D-E, Extended Data Fig. 1C-D). Globally, more than 75% of transcripts contained at least one intron exhibiting post-transcriptional excision (Fig. 1F), whereas over 25% of transcripts contained three or more. Thus, post-transcriptional splicing of polyadenylated, chromatin-associated RNA is a widespread feature of most transcripts.

### Intermediate isoforms are undergoing active processing

Introns that are not excised while the transcript is associated with chromatin are likely removed eventually, as most transcribed pre-mRNA splice junctions are successfully spliced^25^. However, a small subset of post-transcriptionally excised introns (detained introns) are retained in nuclear polyadenylated transcripts, which can be degraded in an exosome-dependent manner^26–29^. To determine whether chromatin-associated intermediate isoforms are targeted for nuclear RNA decay, we depleted the catalytic subunit of the nuclear exosome, EXOSC10 (Extended Data Fig. 1E-F). We observed neither large-scale changes in global intron retention (Extended Data Fig. 1G) nor upregulation of most intermediate isoforms (Extended Data Fig. 1H, Supplemental Table 2), indicating that these transcripts are not frequently targeted for exosome-dependent RNA decay.

As an orthogonal approach to confirm that intermediate isoforms represent RNAs that are actively undergoing processing, we took advantage of direct RNA nanopore sequencing estimates of the lengths of poly(A) tails^30^, which are added to the 3’ ends of RNA after co-transcriptional cleavage of the nascent RNA. We observed that poly(A) tails grew as splicing progressed, both globally and for individual genes (Fig. 1G-H, Extended Data Fig. 1I). These observations demonstrate that splicing and polyadenylation occur in parallel and emphasize that intermediate isoform reads represent RNAs engaged in active processing.

### Splicing order is defined across multiple introns in pre-mRNAs

In many genes with more than one post-transcriptionally excised intron, certain introns were present only if another intron was also present. For example, in *DDX39A*, intron 8 was almost exclusively present when intron 6 was also present (Fig. 2A). These observations suggest that removal of some introns occurs only after other introns have been excised from the transcript and that post-transcriptional splicing follows a defined order.

**Figure 2.**
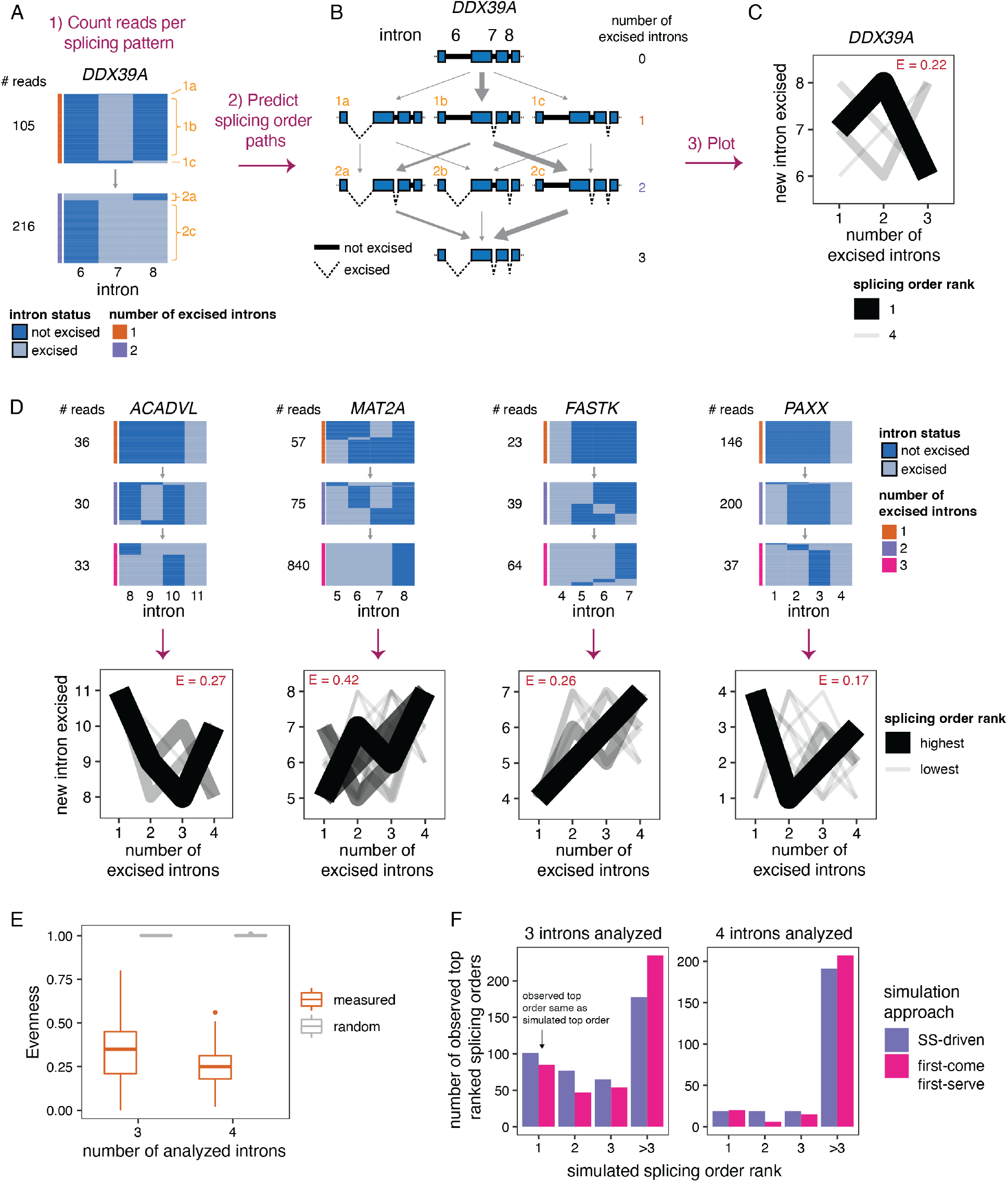
Post-transcriptional splicing follows a defined order. A) Representation of all reads mapping to introns 6 to 8 of *DDX39A*. Each column represents one intron, with light blue showing present introns while dark blue indicates excised introns. The horizontal blocks represent the proportion of reads for each intermediate isoform, with the total number of reads shown on the left. The side color bar shows the number of excised introns in the intron group for each read. Orange numbers identify each intermediate isoform. B) Schematic of splicing order analysis of introns 6 to 8 of *DDX39A*. Each possible intermediate isoform, from fully unspliced to fully spliced, is depicted according to the number of excised introns. Arrows indicate the possible paths to go from one intermediate isoform to another. Excised introns are represented by dotted lines, while present introns are shown as full thicker lines. Intermediate isoforms are numbered in orange as in A). C) Splicing order plot for *DDX39A* introns 6 to 8. The thickness and opacity of the lines are proportional to the frequency at which each splicing order is used, with the top ranked order per intron group set to the maximum thickness and opacity. The evenness (E) is shown in red font. D) Top: Representation of all reads mapping to four example intron groups, as in A). Bottom: Splicing order plots for 4 example intron groups, shown as in C). E) Evenness of splicing order across all analyzed intron groups, separated by the number of introns in each group (3 or 4). Evenness is compared for measured (orange) or random (grey) splicing order. Boxplots elements are shown as follows: center line, median; box limits, upper and lower quartiles; whiskers, 1.5x interquartile range; points, outliers. F) Simulation of splicing order ranks based on splice site score^53^ (SS-driven) or transcription order (first-come first-serve). The top ranked splicing order observed in the data was extracted for each intron group and compared to the splicing order ranks obtained by simulation (x-axis). The majority of observed top ranked splicing orders have a rank of above 3 in the simulations.

To perform a broader analysis of post-transcriptional splicing order, we identified sets of three or four introns (“intron groups”) undergoing post-transcriptional splicing within a single transcript. We developed an algorithm to predict the most frequent orders in which groups of introns are removed to go from fully unspliced to fully spliced. Our method calculates the frequency of each pre-mRNA intermediate isoform, assembles the possible splicing orders, predicts the relative flux through each one, and assigns them a score so that all orders can be ranked (Fig. 2A-C). For example, for introns 6 to 8 in *DDX39A*, the relative frequencies of intermediate isoforms suggested that intron 7 is usually excised first, followed by intron 8 and finally intron 6 (Fig. 2A-C). Indeed, ranking the splicing order scores revealed that of the six possible paths, splicing followed one order predominantly, with a couple of other orders used much more rarely. We observed similar results for groups of four introns in *ACADVL, MAT2A, FASTK* and *PAXX* (Fig. 2D), in which one or a few orders predominated relative to the 24 possible paths. Direct RNA nanopore sequencing read lengths and throughput restrictions limited a global analysis to shorter introns in highly expressed genes (Extended Data Fig. 2B). Nevertheless, applying this approach to the entire dataset, we obtained reproducible splicing orders for 669 distinct intron groups in 325 genes (Extended Data Fig. 2C, Supplemental Table 3).

To quantify the extent to which splicing order is defined in each intron group, we computed a diversity measure, evenness (E), based on the Shannon diversity index^31,32^. A high evenness value (close to 1) indicates that more splicing orders are being used and that they tend to be used at a similar frequency, while lower values indicate that a smaller number of orders are preferred. We found that evenness values from observed splicing orders were almost all less than 0.5, with a median close to 0.25 or 0.35 for groups of three or four introns, respectively (Fig. 2C-E, Extended Data Fig. 2D, Supplemental Table 4). By contrast, the evenness values for simulated random intermediate isoforms were near 1. The highest evenness values in our dataset were 0.56 (four introns) and 0.8 (three introns); in these cases, more than one order was used with high frequency (Extended Data Fig. 2E); however, this still represented only a subset of the possible splicing orders. Accordingly, splicing order in the top path per intron group was consistent with the splicing index from short-read sequencing of chromatin-associated RNA^14^ (Extended Data Fig. 2F). These splicing orders could not be explained by the order in which the introns are transcribed or by their splice site strengths (Fig. 2F). Together, these findings indicate that multi-intron post-transcriptional splicing converges towards one or a few predominant orders per intron group and that it is predetermined rather than stochastic.

### Multi-intron splicing order is fixed during motor neuron differentiation

Introns flanking alternative exons (AS introns) are enriched in post-transcriptionally excised introns^3^, but the order in which these introns are removed relative to other post-transcriptionally excised proximal introns and how this influences AS remain unknown. Changes in AS are abundant during neurogenesis^33^. In particular, several splicing factors are essential for proper motor neuron development and function in mice, and their absence leads to widespread AS changes^34–36^. Thus, we sought to investigate the interplay between AS and splicing order during the differentiation of human induced pluripotent stem cells (iPSC) to spinal motor neurons (sMN)^37^. We collected cells at days 4, 9, and 14 of differentiation, representing neural progenitors, sMN progenitors, and mature sMNs, respectively, and performed direct RNA sequencing of poly(A)-selected chromatin-associated RNA (Fig. 3A, Extended Data Fig. 3A-D). As observed in K562 cells, splicing order was defined at each timepoint, with median evenness values less than 0.4 (Extended Data Fig. 4A). We computed splicing order for all intron groups that were commonly expressed between the three differentiation timepoints and did not include splice junctions that were differentially spliced during differentiation. The majority of intron groups used the same top splicing order in each cell type, revealing that splicing order is largely conserved across differentiation (Extended Data Fig. 4B). Remarkably, this was also the case when comparing splicing order between K562 cells and the three sMN differentiation timepoints (Fig. 3B, Supplemental Table 5). Thus, splicing order is largely conserved between a cancer cell line and non-cancer cell types and throughout sMN differentiation, indicating that splicing order is a fixed phenomenon with low inter-cell type variability.

**Figure 3.**
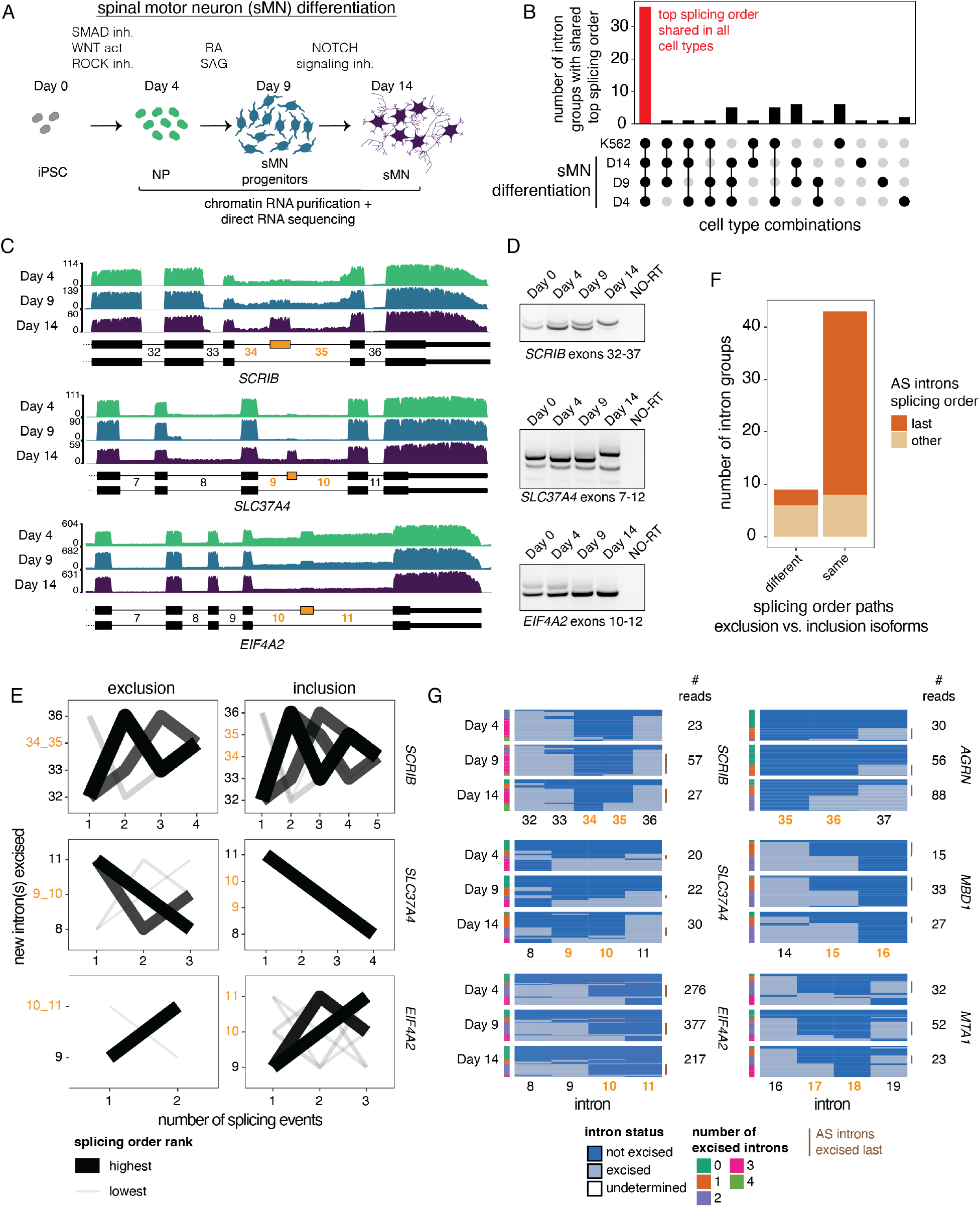
Introns flanking alternative exons (AS introns) have defined splicing order during motor neuron differentiation. A) Schematic of the differentiation of spinal motor neurons (sMN) from human induced pluripotent stem cells (iPSC). Samples were collected at days 4, 9 and 14 of differentiation for cellular fractionation, isolation of chromatin-associated RNA, and direct RNA sequencing. NP: neural progenitors, inh: inhibitor, act: activator, RA: retinoic acid, SAG: Smoothened agonist. Schematic was adapted from^37^ and made with images from https://smart.servier.com/. B) UpSet plot showing the overlap in top splicing order between K562 cells and the three sMN differentiation timepoints. The y-axis indicates the number of intron groups with the same top ranked splicing order in the cell type combinations shown on the x-axis. A filled black circle indicates that a given cell type is included in the combination. C) Nanopore direct RNA sequencing coverage tracks for three example alternative cassette exons with differential inclusion during sMN differentiation. Gene structures are shown for each gene, with introns as lines and exons as rectangles. Alternative exons are depicted in yellow. Introns are numbered and AS introns are shown in yellow font. D) Validation of AS events by RT-PCR of total RNA. The region amplified by PCR is indicated below each gel. NO-RT: no reverse transcriptase control. E) Splicing order plots for three intron groups that contain AS introns, shown separately for inclusion and exclusion of the alternative exon. All timepoints were combined. AS introns are shown in yellow font. The thickness and opacity of the lines are proportional to the frequency at which each order is used, with the top ranked order per intron group set to the maximum thickness and opacity. F) Number of intron groups in which the introns are excised in the same order or in a different order in the top ranked splicing orders between isoforms that show inclusion or exclusion of an alternative exon. Intron groups are further separated by whether the AS introns are removed last in the top ranked splicing order or spliced earlier. G) Representation of all reads mapping to six example intron groups, separated by timepoint. Each column represents one intron, with light blue showing excised introns while dark blue indicates present introns. The horizontal blocks represent the proportion of reads for each intermediate isoform, with the total number of reads shown on the right. The side color bar shows the number of excised introns in the intron group for each read. AS introns are shown in yellow font. Reads in which the AS introns are not removed and the neighboring introns are excised, indicating that the AS introns are removed last, are shown by a brown vertical line.

Our data revealed that AS remodeling is widespread during human sMN differentiation: we identified 624 distinct introns that neighbor alternative cassette exons or alternative splice sites (Supplemental Table 6, Fig. 3C-D, Extended Data Fig. 3E) with inclusion levels that changed significantly between at least two timepoints. To investigate connections between AS and splicing order, we defined intron groups consisting of the AS introns surrounding an alternative exon (Fig. 3C), as well as up to three upstream and downstream introns. Because splicing orders were stable across differentiation, we combined reads from all timepoints and computed splicing orders separately for the reads that included or excluded the alternative exon (Supplemental Table 7). Interestingly, whether or not the alternative exon was included, introns were generally removed in the same order (83% of intron groups, Fig. 3E-F). Furthermore, in isoforms that followed the same splicing order, the introns flanking the alternative exon were most often (81%) excised last in the top-ranked splicing order (Fig. 3E-F). For example, introns 34 and 35 in *SCRIB*, a gene required for zebrafish motor neuron migration^38^, and introns 9 and 10 in *EIF4A2* were removed after their neighboring introns whether they were excised together with the cassette exon or separately, leaving the exon in the mature transcript (Fig. 3E). In a smaller number of intron groups (e.g. *SLC37A4*, Fig. 3E-F), AS introns were removed earlier in the top ranked order, but remained at the same positions regardless of inclusion or exclusion.

These results could also be recapitulated at each timepoint, where reads supporting AS introns being excised last were more frequent than reads supporting other splicing orders (Fig. 3G, Extended Data Fig. 4C). These results indicate that, for the most part, splicing order is not differentially regulated between different isoforms, but rather programmed for the AS introns being removed later, further emphasizing the fixed nature of splicing order.

### U2 snRNA depletion modifies the splicing order of many transcripts

Given that splicing order is largely conserved across cell types and among alternative isoforms, we sought to investigate the consequences of disrupting the order of intron removal. We previously reported that binding of the U2 snRNP is correlated with splicing order of intron pairs^14^, so we depleted U2 snRNA in HeLa cells in order to ask whether U2 snRNP levels control splicing order. We used an antisense oligonucleotide (ASO)^22^ to reduce U2 snRNA levels in HeLa cells to ∼25–50% of control levels (Fig. 4A). Importantly, we observed no difference in RNA polymerase II promoter-proximal pausing index between the control and U2 snRNA knockdown (KD) (Extended Data Fig. 5A), indicating that this modest depletion of U2 snRNA did not affect transcription dynamics in the same manner as a strong inhibition of the U2 snRNP by the small molecule pladienolide B^39^. Global co-transcriptional splicing kinetics and splicing order were not affected^14^ (Extended Data Fig. 5B-C). However, short-read RNA-seq revealed numerous intron retention and exon skipping events (Fig. 4B-C, Extended Data Fig. 5D), as previously reported^22^. Interestingly, we observed that genes with retained introns (RIs) were more likely to also contain skipped exons (SEs) (Extended Data Fig. 5E). Direct mRNA nanopore sequencing showed that for most genes with RIs and SEs upon U2 snRNA KD, these two events frequently occurred together on the same RNA molecules (Fig. 4D-E, Extended Data Fig. 5F).

**Figure 4.**
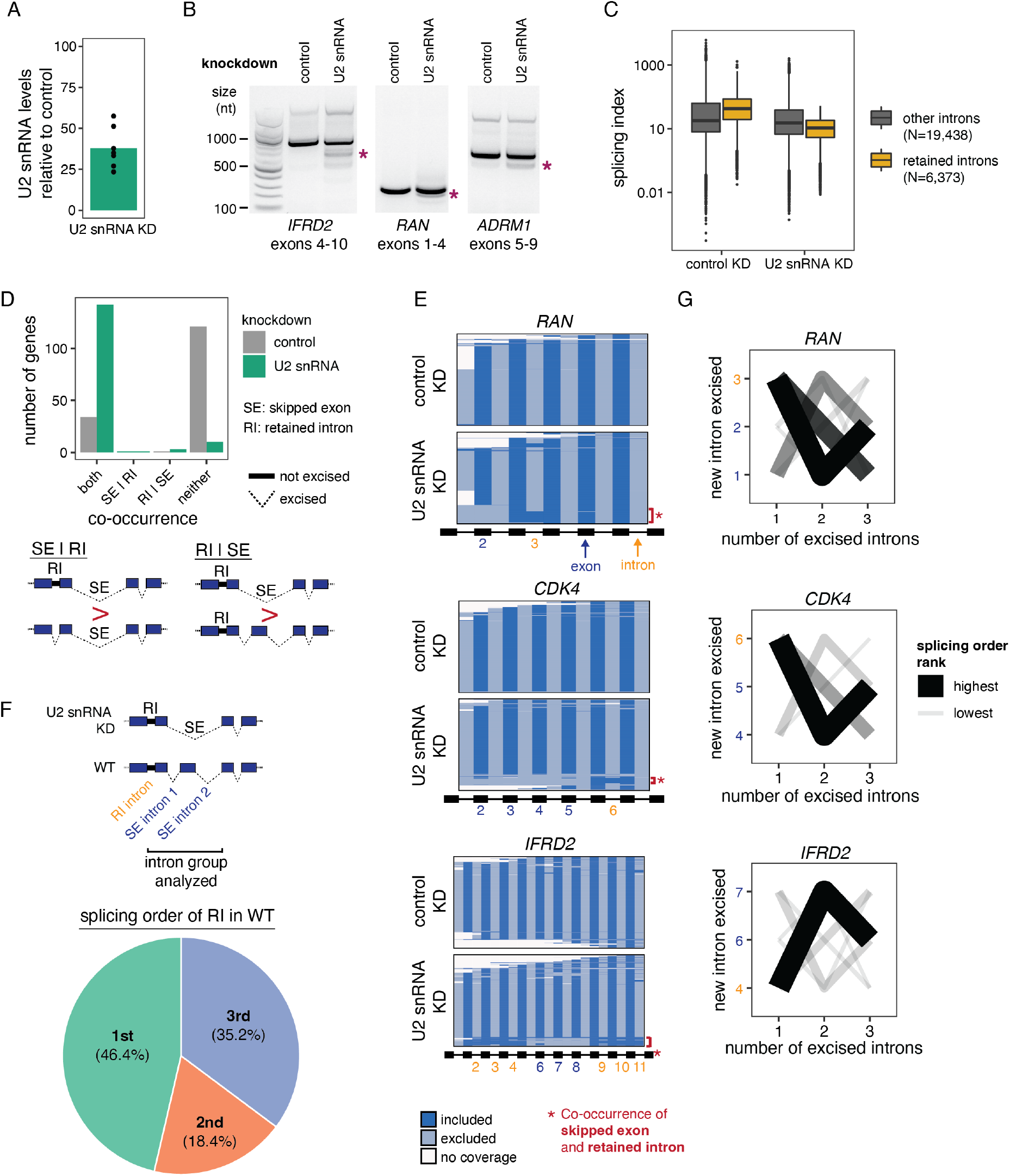
Splicing order changes upon U2 snRNA-mediated exon skipping. A) qRT-PCR showing reduced U2 snRNA levels relative to control after a 48 hour KD with 200 pmoles of an ASO targeting U2 snRNA. U2 snRNA levels were normalized to 5S rRNA and 7SL RNA levels using the ΔΔCt method. B) Agarose gel electrophoresis of RT-PCR products showing alternative splicing events upon U2 snRNA KD. Asterisks indicate exon skipping events. C) Distribution of total RNA splicing index for introns that are normally excised (“other”) or that are retained upon U2 snRNA KD, in the control and U2 snRNA KD conditions. The average across biological triplicates was used for each intron. The number of introns (N) in each category is indicated. Normally excised and retained introns were compared using a two-sided Wilcoxon test (p-value < 10^-100^). Boxplots elements are shown as follows: center line, median; box limits, upper and lower quartiles; whiskers, 1.5x interquartile range; points, outliers. D) Number of genes showing co-occurrence of skipped exons (SE) and retained introns (RI). “Both” means that the SE occurs more significantly when the RI is present and the RI occurs more significantly when the SE is present. “SE | RI”, where the vertical bar is the symbol for “given”, means that the SE occurs more significantly given the presence of the RI, while “RI | SE” indicates that the RI occurs more significantly given the presence of the SE. “Neither” indicates cases in which the SE and the RI occur at similar frequencies together and separately. Schematics depicting “SE | RI” and “RI | SE” are shown below the plot. E) Representation of all reads mapping to *RAN*, *CKD4* and *IFRD2* upon direct RNA sequencing of poly(A)-selected RNA from control or U2 snRNA KD. Each line represents one read and each column represents an intron or an exon. Light blue represents exclusion (excised intron or skipped exon), while dark blue depicts inclusion (retained intron or included exon) and white indicates an absence of coverage. The gene structure is displayed at the bottom (not at scale), with introns shown as lines and exons as rectangles. RIs and SEs in the U2 snRNA KD are shown as yellow or dark blue numbers, respectively. Red brackets with asterisks highlight reads in which co-occurrence of SE(s) and RI(s) is observed. F) Splicing order in wild-type (WT) K562 cells of introns involved in co-occurring SEs and RIs upon U2 snRNA KD. The top schematic shows how the intron groups were defined. The pie chart shows the order in which the RI is removed within the intron group. G) Splicing order plots for *RAN*, *CDK4* and *IFRD2* in WT K562 cells, for groups containing introns involved in exon skipping and intron retention upon U2 snRNA KD. On the y-axis, introns involved in intron retention are shown in orange and those involved in exon skipping are shown in dark blue. The thickness and opacity of the lines are proportional to the frequency at which each order is used, with the top ranked order per intron group set to the maximum thickness and opacity.

Globally, introns retained upon U2 snRNA KD had a higher splicing index in control cells than unaffected introns (Fig. 4C, Extended Data Fig. 5D), suggesting that introns that are sensitive to U2 snRNA KD are normally excised more rapidly than other introns and that U2 snRNA KD disrupts splicing order in the affected transcripts. Consistently, when inspecting the splicing orders that we annotated deeply in K562 cells, introns involved in RI events were largely removed first or second, with the introns flanking the SEs excised later (Fig. 4F-G, Supplemental Table 9). Thus, U2-mediated AS is associated with splicing order changes in which introns flanking SEs tend to be removed earlier than normal, whereas excision of RIs, which normally occurs earlier, is delayed or inhibited. Our findings indicate that splicing order is regulated by the spliceosome itself and underscore the importance of a fixed splicing order for controlling the cis-element landscape during splicing regulation.

### Early removal of specific introns in *IFRD2* induces retention of proximal introns

To investigate how splicing order changes lead to intron retention, we analyzed splicing of *IFRD2* transcripts, in which skipping of exons 6-8 upon U2 snRNA KD was almost always associated with retention of three flanking introns on each side (introns 2–4 and 9–11) (Fig. 4E, Extended Data Fig. 6A). Using a targeted PCR-based approach to study splicing order of introns 4–9 in this gene with greater detail (Extended Data Fig. 6B), we observed that in normal conditions, introns 4 and 9 were usually removed first and introns 6–8 were removed later (Extended Data Fig. 6C-D). However, when U2 snRNA-mediated exon skipping occurred, removing introns 5–8 together, splicing order tended to be reversed, with introns 5–8 excised before introns 4 and 9 (Extended Data Fig. 6C-D).

To understand how splicing order determines the final isoform in *IFRD2*, we used ASOs to artificially alter the splicing order of *IFRD2* by blocking the 3’SS of introns 4, 5, and 9 individually or in combination (Fig. 5A, Extended Data Fig. 7). When we disrupted the predominant splicing order for *IFRD2*, in which intron 9 is removed first (Extended Data Fig. 6D), by blocking the 3’SS of intron 9, exon skipping did not co-occur (Fig. 5B). However, impeding the second and third most frequent splicing orders by simultaneously targeting the 3’ SSs of introns 4 and 5 led to a large exon skipping event (removal of introns 4–8) that was associated with retention of intron 9 (Fig. 5B-C). Intron 9 was rarely retained on its own (Extended Data Figure 7C), suggesting that the SE event drives retention of intron 9. These AS patterns were similar to what we observed in the U2 snRNA KD and indicate that alterations in splicing order of upstream introns are sufficient to elicit retention of intron 9.

**Figure 5.**
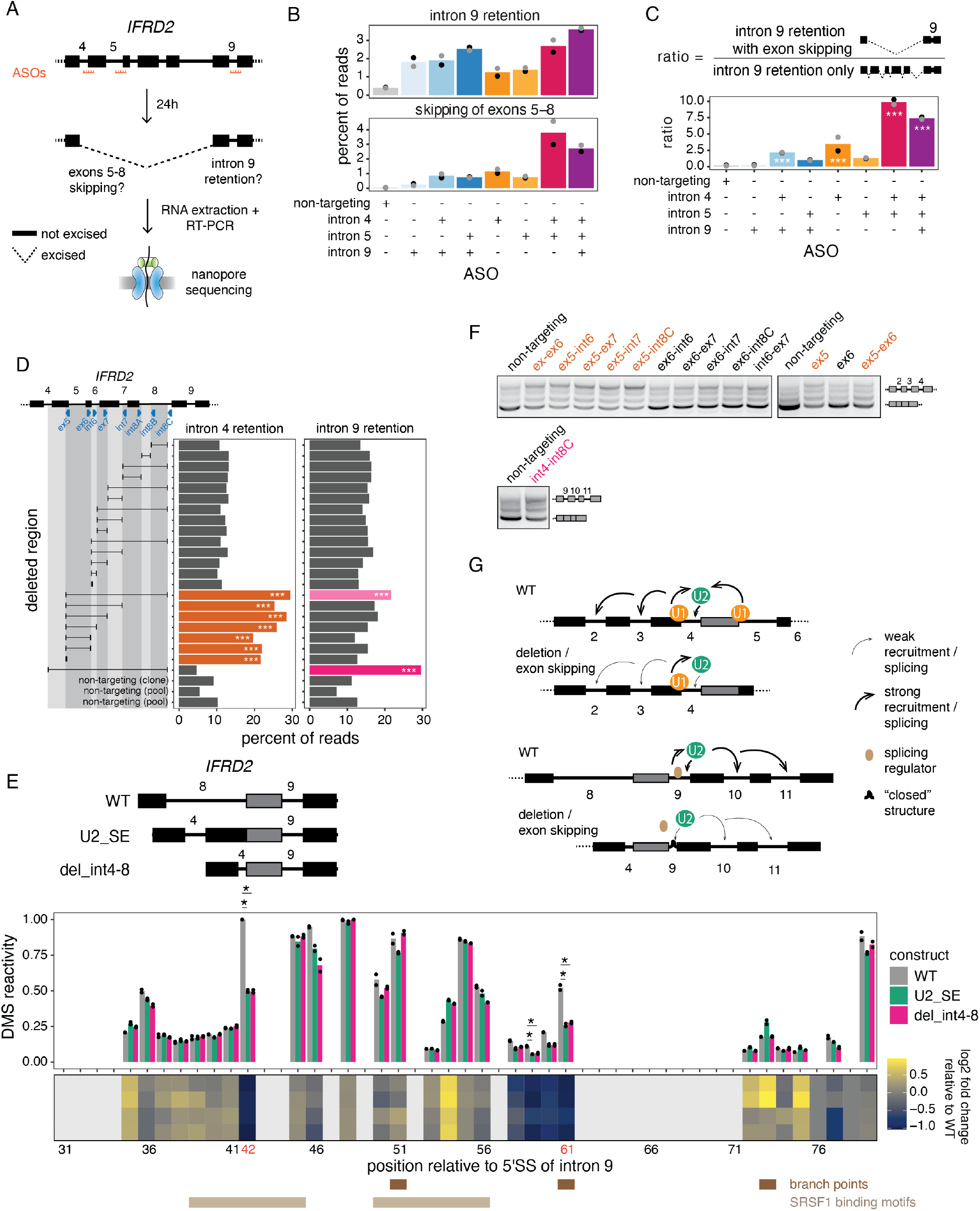
Perturbation of one intron disrupts excision of proximal introns. A) Schematic of the experimental design. Antisense oligonucleotides (ASOs) targeting the 3’SS of introns 4, 5 and 9 of *IFRD2* were transfected individually or in combinations in HeLa cells for 24 hours. Total RNA was extracted and underwent reverse transcription and amplification of *IFRD2* exons 4 to 10. The resulting amplicons were sequenced by nanopore. The gene structure of *IFRD2* is shown at the top, with introns represented by lines and exons by rectangles. The two AS events that are measured in B) are depicted in the middle panel. B) Levels of intron 9 retention or exons 5-8 skipping upon treatment with ASO(s) targeting *IFRD2* or a non-targeting control. Dots indicate biological replicates. The x-axis shows which introns were targeted by the ASOs. C) Ratio of the number of reads with intron 9 retention and exons 5-8 skipping relative to the number of reads with intron 9 retention only upon treatment with ASO(s) targeting *IFRD2* or a non-targeting control. Dots indicate biological replicates. The x-axis shows which introns were targeted by the ASOs. A one-sided binomial test was used to determine whether intron 9 retention occurs more frequently with exon skipping than expected by chance. ***: p-value < 0.001 and ratio >= 2. D) Deletion tiling of *IFRD2* using CRISPR-Cas9 editing and dual sgRNAs. Each sgRNA is represented as a blue triangle below the schematic of the *IFRD2* gene. For each individual sgRNA or sgRNA pair, the resulting deletion is shown as a black horizontal line delimited by vertical lines aligned to the sgRNA(s) used. cDNA-PCR nanopore sequencing of chromatin-associated RNA was used to measure the levels of intron 4 and intron 9 splicing, which are shown as the percent of total reads for each deletion. The names of the sgRNAs are indicated next to deletions that show increased intron retention (“ex”: exon, “int”: intron). Pools of edited cells were used for this experiment, except for the largest deletion (int4-int8C), which was found in a clonal cell line derived from editing with the sgRNA int8C only. Non-targeting controls from pools of cells and a clonal cell line are also shown. A one-sided binomial test was used to compare the frequency of intron retention in each deletion relative to the mean frequency across non-targeting controls. ***: p-value < 0.001 and fold change of percent of reads with intron retention relative to non-targeting controls >= 2. E) Top: *In vitro* transcribed RNAs used for *in vitro* DMS-MaPseq of *IFRD2*. U2_SE imitates the exon skipping event observed upon U2 snRNA KD and del_int4-8 imitates the largest CRISPR-induced deletion. All three RNAs have in common exons 9 to 10 and differ by the upstream region. Middle: DMS-MaPseq reactivity for nucleotides 30 to 80 of *IFRD2* intron 9. Individual dots represent two biological replicates. Bottom: Heatmap showing the log2 fold change of DMS reactivity in U2_SE or del_int4-8 relative to WT. Positions corresponding to G’s and T’s, which do not react with DMS, are shown in light grey. Positions where the largest fold change is observed in both mutated constructs are shown in orange font. Predicted branch points^41,54^ are shown as brown rectangles and the two predicted SRSF1 binding sites (ESEFinder^42^) with the highest scores are shown as beige rectangles. Reactivities between constructs were compared with a two-sided Student’s t-test. *: p-value < 0.05 and fold change relative to WT >= 2. F) Agarose gel electrophoresis showing the RT-PCR products from amplification of introns 2 to 4 of *IFRD2* for a subset of the deletions shown in D) and from amplification of introns 9 to 11 for the largest deletion (int4-int8C, pink font). Deletions involving the end of exon 5 are shown in orange font. The identity of each RT-PCR product is shown on the right of the gel. G) Working models for the two mechanisms regulating splicing of introns 4 and 9. Top: U1 snRNP bound to the unspliced 5’SS of intron 5 helps to recruit U2 snRNP to the upstream BPS of intron 4. If intron 5 is already excised or if the 5’SS is deleted, U2 snRNP is less efficiently recruited to intron 4, which is more frequently retained together with introns 2 and 3. Bottom: in normal conditions, intron 9 is in an open structure that is accessible to the U2 snRNP and to splicing regulators (e.g. SRSF1) that may help to recruit the U2 snRNP. When exon 4 is in closer proximity to intron 9 due to exon skipping or a deletion, the structure of intron 9 is more closed, leading to decreased binding of U2 snRNP and/or splicing regulators and to increased intron 9 retention, which in turns promotes retention of introns 10 and 11.

### Distinct cis-regulatory mechanisms impact splicing fidelity after splicing order changes in IFRD2

We postulated that delayed removal of *IFRD2* introns 4 and 9 upon U2 snRNA KD was due to the early removal of cis-elements within the region spanning introns 5 through 8, which are necessary to stimulate excision of introns 4 and 9. To test this hypothesis, we used CRISPR-Cas9 and dual sgRNAs to introduce overlapping deletions spanning this entire region (Fig. 5D, Extended Data Fig. 8A-B). Interestingly, deleting as few as 7 nt near the 5’SS of intron 5 was sufficient to increase retention of intron 4 (Fig. 5D), indicating that an intact and unspliced 5’SS in the downstream intron is critical for proper excision of intron 4. By contrast, intron 9 retention only occurred when the entire region of interest was deleted (Fig. 5D), suggesting that there is no single sequence in the upstream introns that stimulates intron 9 removal.

To determine whether changes in RNA conformation underlie intron 9 retention, we performed *in vitro* dimethyl sulfate (DMS) mutational profiling with sequencing (DMS-MaPseq)^40^, in which unpaired adenosines and cytosines are modified by DMS and high DMS reactivity is indicative of nucleotides that are frequently single-stranded. To investigate the structure of intron 9 in the presence of different upstream contexts, we *in vitro* transcribed three RNAs: the unspliced WT *IFRD2* transcript from exons 8 to 10 (WT), the transcript produced as a result of U2 snRNA-mediated exon skipping (U2_SE), and the transcript with the largest CRISPR-induced deletion (del_int4-8) (Fig. 5E). DMS reactivity was significantly reduced at one of the predicted intron 9 branch point adenosines^41^ in the U2_SE and del_int4-8 constructs relative to WT (Fig. 5E, Extended Data Fig. 9), suggesting reduced availability for U2 snRNA binding. Moreover, a position in intron 9 that was highly reactive in the WT, indicating that it is normally unpaired, exhibited a substantial decrease in reactivity in both alternative constructs (Fig. 5E, Extended Data Fig. 9). This position overlaps with a predicted splicing enhancer bound by the splicing factor SRSF1 (ESEFinder^42^), and its high reactivity in the WT construct is consistent with the reported preference of SRSF1 for binding single-stranded RNA^43–45^. Thus, our data suggest that intron 9 is more open at certain positions in the WT sequence, but when it is in closer proximity to exon 4 upon upstream exon skipping or deletion, structural rearrangements render it less accessible and thus more protected from splicing factors than in the WT RNA. Together, these results demonstrate that two different mechanisms regulate excision of introns 4 and 9. In both cases, however, removing introns that are usually excised later, either through splicing or a deletion, induced retention of at least one of the two introns that are normally excised earlier.

Finally, we asked whether intron retention is perpetuated further down the transcript, as observed upon U2 snRNA KD (Fig. 4D, Extended Data Fig. 5A). Remarkably, deletions that induced intron 4 retention also exhibited higher levels of retention of introns 2 and 3 (Fig. 5F). Similarly, the large deletion that increased retention of intron 9 also led to retention of introns 10 and 11 (Fig. 5F). Together, our findings indicate a strong interdependency in removal of introns in *IFRD2* (Fig. 5G), supporting the notion that splicing is coordinated between neighboring introns^14^ and highlighting the importance of maintaining proper splicing order in order to generate a fully spliced mature mRNA.

## Discussion

Here, we combined direct RNA nanopore sequencing with a novel algorithm to show that multi-intron splicing order is predetermined and conserved across different cell types. We found that splicing order can be perturbed through modulation of U2 snRNA levels, direct physical inhibition of a SS, or genomic deletion, resulting in long-range disruptions in cis that affect many introns along the transcript. These results underscore the coordinated nature of the many splicing events that occur along a transcript and demonstrate that perturbing the excision of a single intron can have far-reaching and long-lasting consequences on the final isoform.

Our findings expand on previous studies showing that splicing order is defined for pairs of consecutive introns transcriptome-wide^13,14^, as well as with single-gene experiments demonstrating that multiple introns within a given transcript are predominantly excised in one or a few orders^15–17^. We observed that introns flanking alternative exons are most often removed last, after their neighbors, suggesting that splicing order is set in order to predispose these introns to be excised last no matter the final isoform. This may allow more time for splicing regulators to bind to these introns and steer the splice site choice, or may instead represent a reservoir of almost mature transcripts that can be fully spliced into either isoform depending on cellular needs.

Due to the current read length of nanopore sequencing, we were limited to analyzing groups of three or four proximal introns that are shorter than the genomic average. Consequently, our findings may not hold true for longer introns. Yet, recent work showed that longer introns (>10 kb) are frequently removed in smaller chunks through stochastic recursive splicing, whereas shorter introns (<1.5kb) are mostly excised as complete units^46^. It is reasonable to speculate that removal of shorter canonical or recursive introns follows a particularly well-defined order, where excision of proximal introns is dependent on one another because of the shorter distance between them. Moreover, due to more limited coverage of direct RNA sequencing, our analyses were restricted to introns in highly expressed genes. Nonetheless, many of these genes accomplish essential functions in gene expression and metabolism, highlighting the importance of understanding how splicing proceeds across these transcripts.

The high degree of consistency of splicing order across cell types and isoforms suggests that it is robustly defined. Previous analyses of human intron pairs showed that intron length, SS sequence, GC content, sequence motifs, and RNA-binding protein (RBP) binding are associated with splicing order, but that these features cannot explain a considerable fraction of the variance, suggesting that additional factors are at play^13,14,47^. We found that moderate depletion of U2 snRNA levels changed splicing order in many genes. Interestingly, when we reanalyzed nanopore sequencing data from the cerebella of U2 snRNA mutant (*NMF291^-/-^)* mice^48^, we found a similar co-occurrence of RIs and SEs (Extended Data Fig. 5G-H), indicating that splicing order is partially determined by the spliceosome and that splicing fidelity is disrupted when splicing order changes *in vivo*. Our analysis of *IFRD2* splicing order suggests that both cis-elements and secondary structure also play important roles (Fig. 5G), possibly by modulating the landscape of RBPs in introns^49,50^. Identifying the determinants of splicing order on a broader scale will presumably require mapping of RBP binding and secondary structure in an intermediate isoform–specific manner, in which each transient intermediate isoform can be associated with its specific features. Nanopore sequencing was recently used to detect RBP binding sites^51^ and reveal isoform-specific secondary structure of mRNA^52^, suggesting that future experiments combining these approaches and splicing order analyses could uncover the global role of RBPs and RNA structure in the determination of splicing order.

We found that early removal or a small deletion in the 5’SS of an intron is sufficient to induce retention of three upstream introns in *IFRD2*, demonstrating for the first time that cis-elements can have longer-range effects on several proximal introns. These results are consistent with *in vitro* findings showing that U1 snRNP binding to the 5’SS of an intron and the next downstream 5’SS exerts synergistic effects on U2 snRNP recruitment to the intron BPS and increases splicing efficiency^24^. This observation also adds to the evidence that excision of neighboring introns is coordinated in living cells^14^. The distance between introns 2 and 5 of *IFRD2* is relatively short (∼800 nt), and this cross-intron coordination may be a feature of shorter introns. Alternatively, the effect could be due to a ripple effect of shorter-range interactions where the 5’SS of intron 5 is essential for intron 4 removal, intron 4 has an element that impacts intron 3, and so on. Further experiments are required to determine how these long-range effects occur and whether the mechanism of splicing regulation in *IFRD2* represents a special case or is widespread.

An understanding of splicing order transcriptome-wide has important implications for human health. The order of intron removal across several introns influences the splicing outcome of disease-causing mutations^15,16^, indicating that knowledge of splicing order could help interpret the impact of genetic variants. Moreover, splicing order can influence the efficiency of ASO-mediated exon skipping, used as a therapeutic strategy^17^. Consequently, taking splicing order into account could help to devise efficient RNA-based therapeutic approaches by ensuring that splicing fidelity is maintained for all introns in a targeted transcript.

---

**Extended Data Figure 1 (related to Figure 1).**
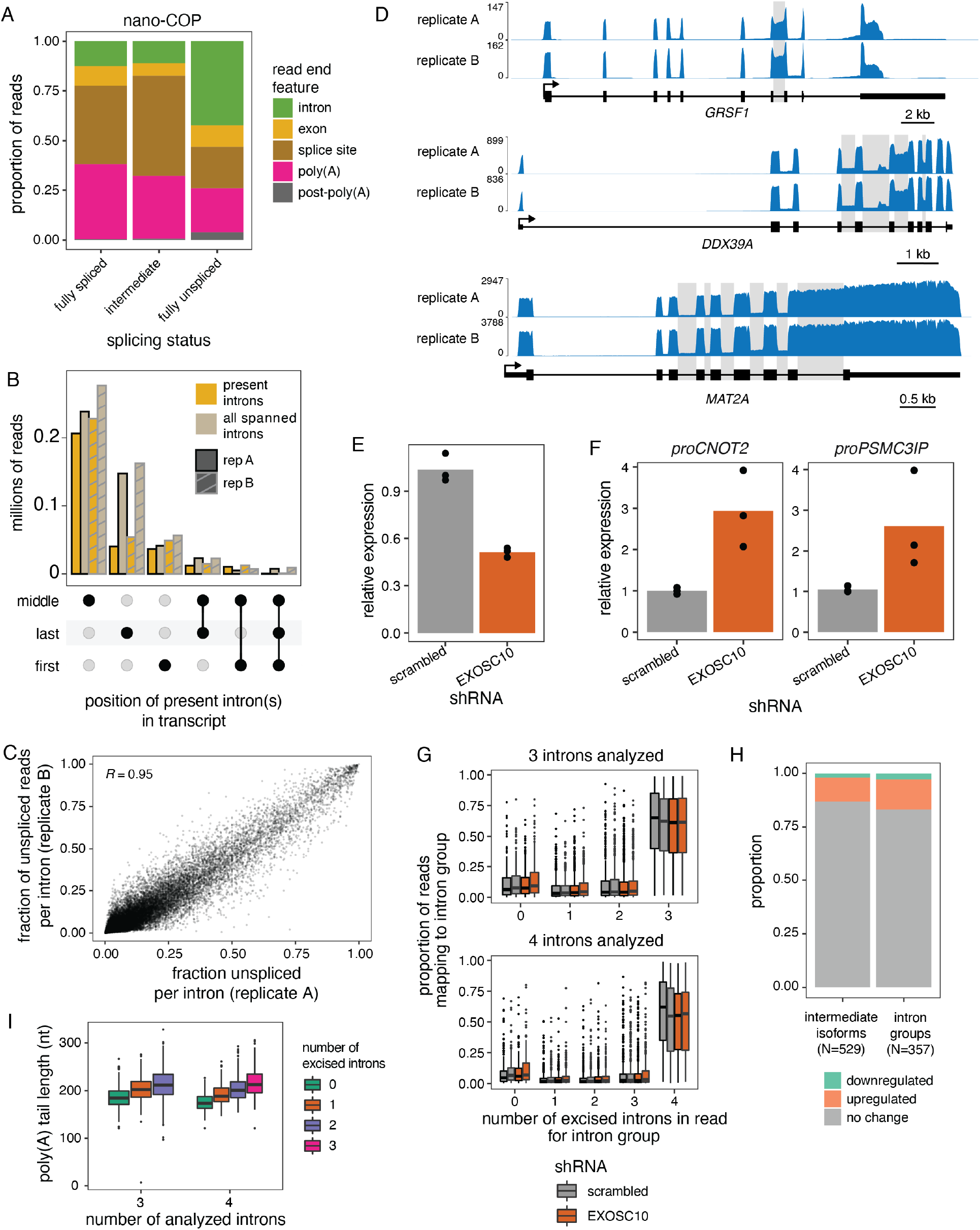
A) Distribution of RNA 3’ends based on read splicing status in nano-COP data from K562 cells. 3’end features are defined as in ^14^. B) UpSet plot showing the position of introns present in partially spliced reads from direct RNA sequencing of poly(A)-selected chromatin-associated RNA. The y-axis axis shows the number of million reads for each combination of intron positions displayed on the x-axis. The number of present introns is compared to the total number of spanned introns (present or excised) in partially spliced reads. “Middle” includes introns that are not the first or last intron in a transcript. Two biological replicates are displayed. C) Scatter plot of the percent of unspliced reads per intron between two biological replicates. Pearson’s correlation coefficient is shown on the plot (p-value < 0.0001, 95% confidence interval 0.948-0.950). Only reads spanning two introns or more are included. D) Coverage tracks from direct RNA sequencing of poly(A)- selected chromatin-associated RNA for three genes displaying various levels of post-transcriptional splicing. Introns with fraction of unspliced reads > 0.1 are shaded in grey. The gene structure is shown for each gene, with rectangles representing exons and lines indicating introns. The arrow indicates the transcription start site. E) *EXOSC10* mRNA levels and F) RNA levels of two promoter upstream transcripts (PROMPT) following shRNA-mediated knockdown (KD) of EXOSC10, as measured by qRT-PCR of total RNA. *EXOSC10* or PROMPT levels were normalized to *ACTB* levels using the ΔΔCt method. Dots represent biological replicates. G) Distribution of the proportion of reads mapping to each intron group used in splicing order analyses (Fig. 2) as a function of the number of excised introns in the read. Biological duplicates for each shRNA treatment are shown side by side. H) Proportion of intermediate isoforms and intron groups that show a significant change (FDR < 0.1 and odds ratio > 1 or < -1) in abundance upon EXOSC10 KD compared to a scrambled control. The total number of isoforms and intron groups is shown below the x-axis. I) Distribution of median poly(A) tail lengths as a function of splicing level. All reads are classified into groups of 3 or 4 introns based on the number of proximal post-transcriptionally excised introns that the read covers. Each intron group must have at least 10 intermediate isoform reads for each number of excised introns in the read. The median poly(A) tail length across reads is calculated for each number of excised introns in each intron group. In G) and I), boxplots elements are shown as follows: center line, median; box limits, upper and lower quartiles; whiskers, 1.5x interquartile range; points, outliers.

**Extended Data Figure 2 (related to Figure 2).**
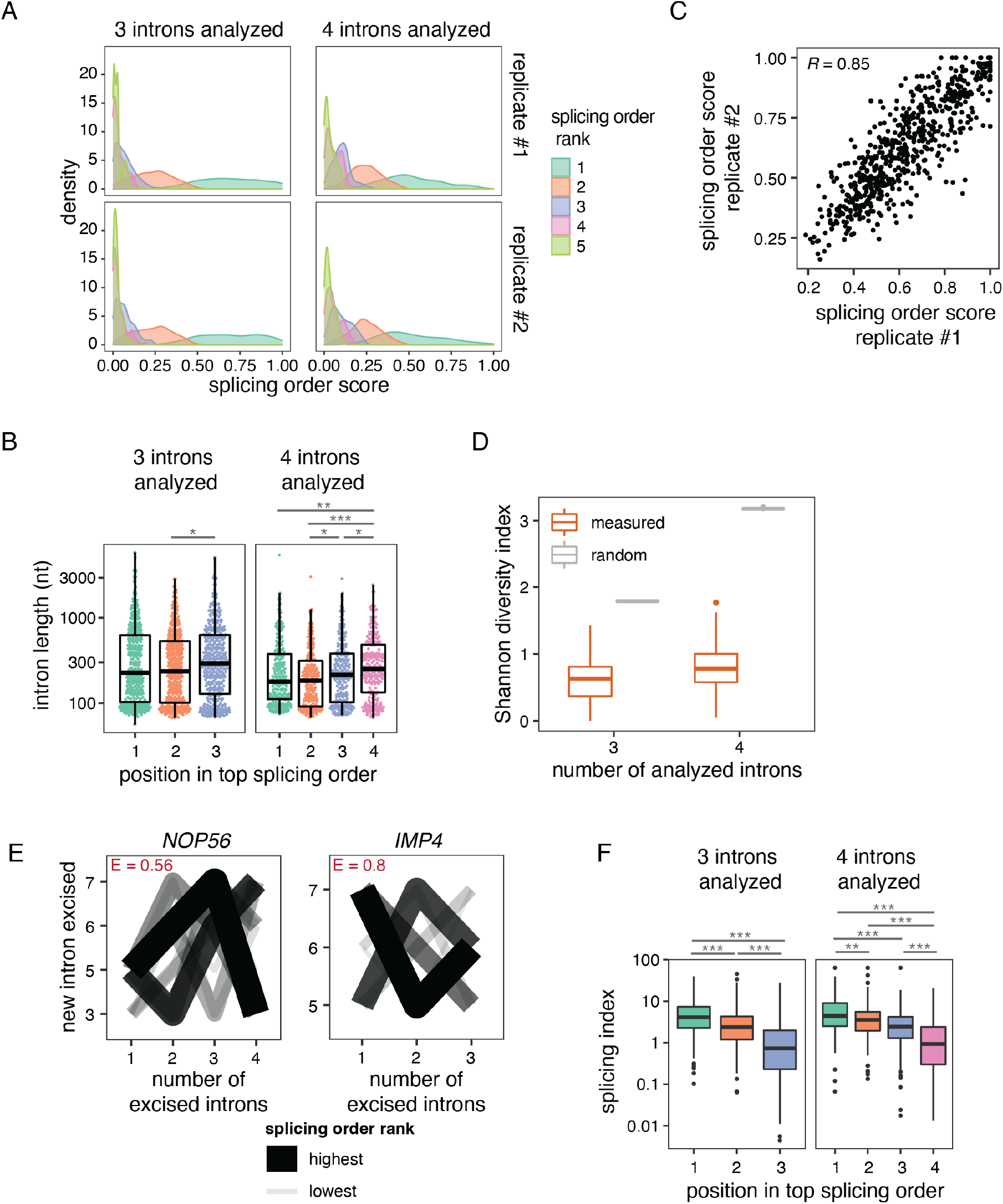
A) Distribution of splicing order scores, colored by splicing order rank and separated by the number of introns in each group (3 or 4). Splicing orders that are ranked from 1 to 5 are included. Two biological replicates are shown. B) Distribution of intron length as a function of the position of introns in the top ranked splicing order for each intron group. C) Correlation in splicing order scores between two biological replicates for top ranked splicing orders. Each dot represents one intron group. Pearson’s correlation coefficient is shown on the plot (p-value < 0.0001, 95% confidence interval 0.83-0.87). D) Shannon diversity index of splicing order across all analyzed intron groups, separated by the number of introns in each group (3 or 4). Shannon diversity index is compared for measured (orange) or random (grey) splicing order. E) Splicing order plots for the intron groups with the highest evenness for groups of 3 or 4 introns, respectively. The thickness and opacity of the lines are proportional to the frequency at which each splicing order is used, with the top ranked order per intron group set to the maximum thickness and opacity. The evenness (E) is shown in red font for each intron group. F) Distribution of splicing index from short-read RNA-seq of rRNA-depleted chromatin-associated RNA, as a function of the position of introns in the top ranked splicing order for each intron group from direct RNA sequencing. The splicing index was calculated using reads spanning intron-exon junctions and defined as 2*spliced_count/(unspliced_count_3’SS + unspliced_count_5’SS). For B) and F), introns were compared using a two-sided Wilcoxon rank-sum test. *: p-value < 0.05, **: p-value < 0.01, ***: p-value < 0.001. In B), D) and F), boxplots elements are shown as follows: center line, median; box limits, upper and lower quartiles; whiskers, 1.5x interquartile range. For D) and F), points represent outliers.

**Extended Data Figure 3 (related to Figure 3).**
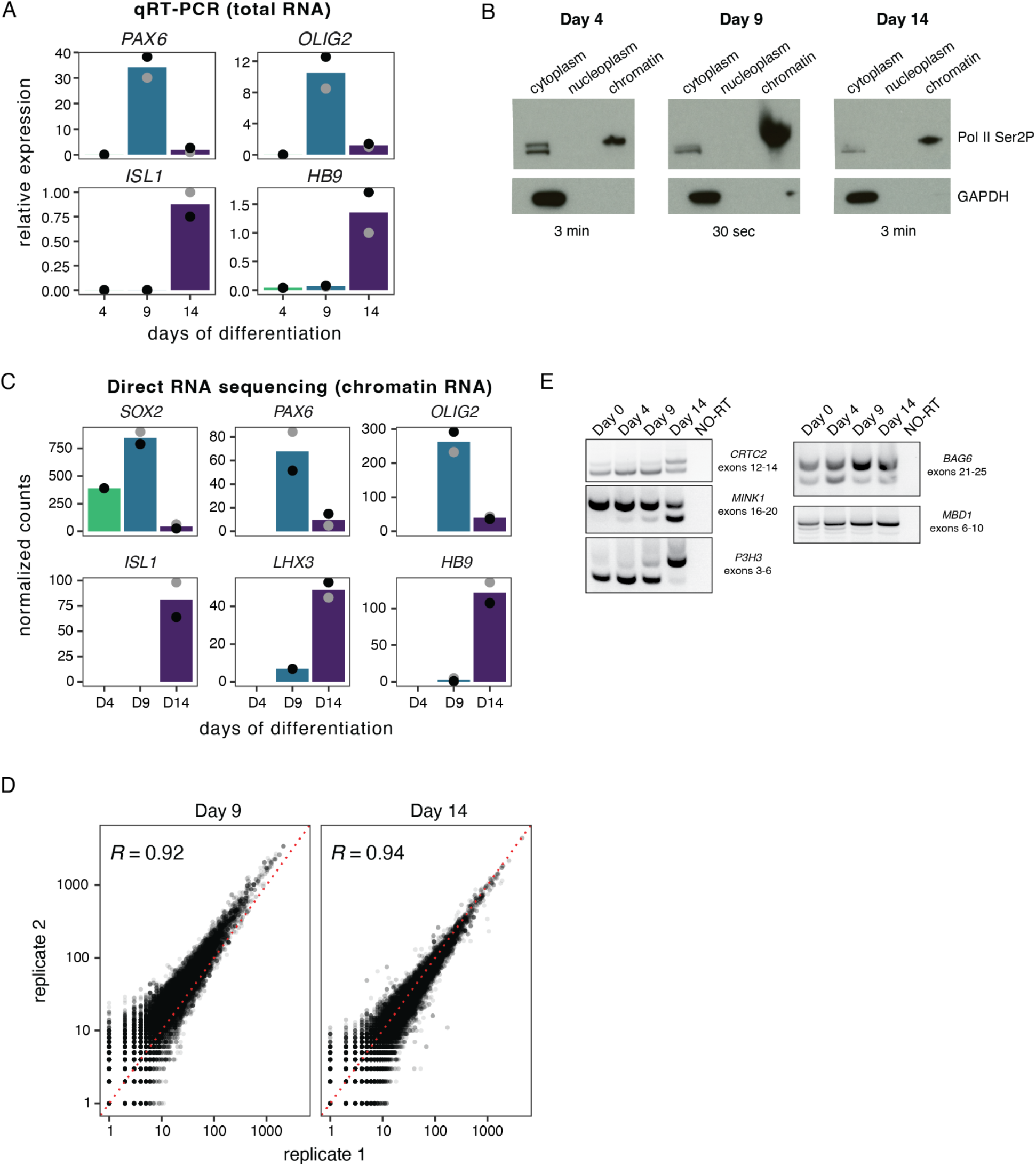
A) Expression of differentiation markers during spinal motor neuron (sMN) differentiation, as measured by qRT-PCR of total RNA. The expression of each differentiation marker was normalized to *ACTB* using the ΔΔCt method. B) Western blot showing the cytoplasm, nucleoplasm and chromatin fractions obtained from cellular differentiation at each sMN differentiation timepoint. GAPDH is exclusively in the cytoplasm, while RNA polymerase II Ser2P is strongly enriched on chromatin. The exposition time used for each blot is noted at the bottom. More cells were used at day 9, resulting in higher abundance of the markers. C) Expression of differentiation markers during sMN differentiation, as measured by direct RNA sequencing of chromatin-associated RNA. The expression of each marker was normalized to sample-specific size factors. D) Scatter plot of reads counts between two biological replicates for direct RNA sequencing of chromatin-associated RNA at days 9 and 14. Pearson’s correlation coefficient is shown on the plots (Day 9: p-value < 0.0001, 95% confidence interval: 0.921-0.924; Day 14: p-value < 0.0001, 95% confidence interval: 0.935-0.938). E) Validation of AS events by RT-PCR of total RNA. The region amplified by PCR is indicated next to each gel. NO-RT: no reverse transcriptase control.

**Extended Data Figure 4 (related to Figure 3).**
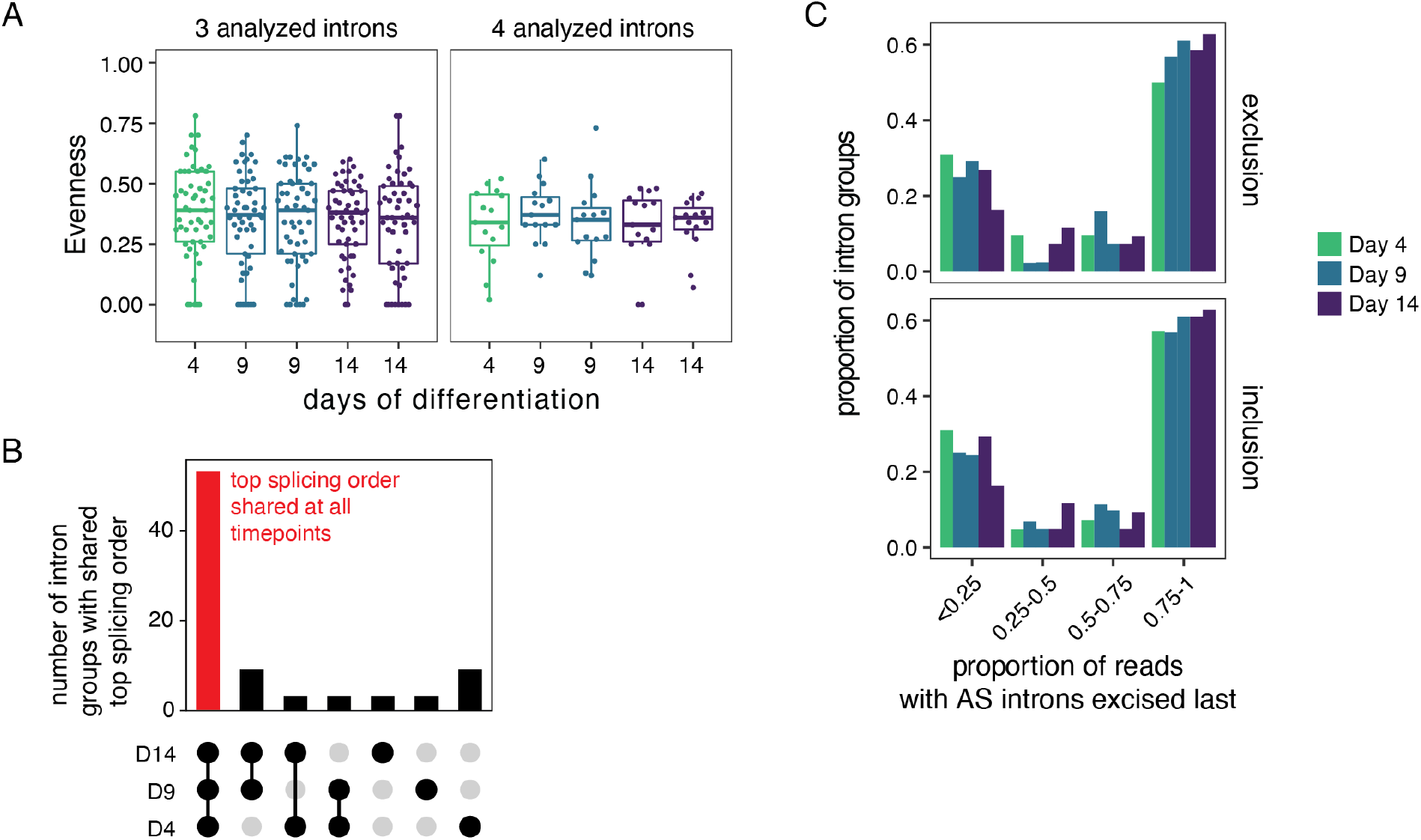
A) Evenness of splicing order for intron groups analyzed at every timepoint of motor neuron differentiation, separated by the number of introns in each group (3 or 4). Biological duplicates are shown for days 9 and 14. Boxplots elements are shown as follows: center line, median; box limits, upper and lower quartiles; whiskers, 1.5x interquartile range; points, intron groups. B) UpSet plot showing the overlap in splicing order between the three sMN differentiation timepoints. The y-axis indicates the number of intron groups with the same top ranked splicing order in the cell type combinations shown on the x-axis. Dark filled circles indicate that a cell type is included in the combination. C) For each timepoint, the proportion of intron groups was plotted as a function of the proportion of reads that show AS introns being excised last (as exemplified by the brown vertical lines in Fig. 3G) relative to all intermediate isoform reads for that intron group. The proportion of reads was calculated separately for those that display inclusion or exclusion of the AS event. Two biological replicates are shown for days 9 and 14, while only one was obtained at day 4.

**Extended Data Figure 5 (related to Figure 4).**
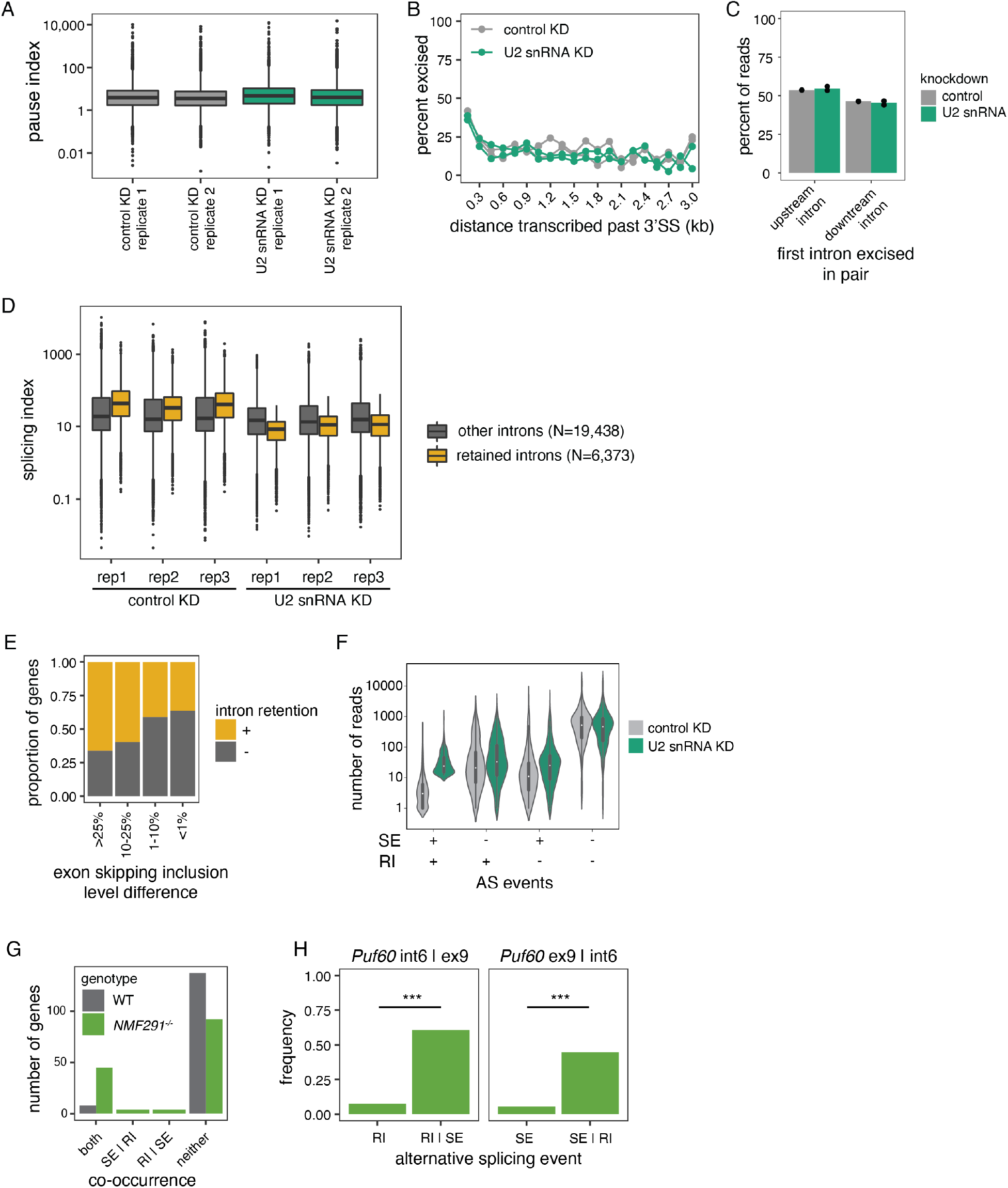
A) Distribution of NET-seq pause index, defined as the RPKM in the promoter region (-30 nt to +300 nt from TSS) divided by the RPKM in the gene body region (+300 nt to end of gene), upon control or U2 snRNA KD. B) Global analysis of the percent of excised introns as a function of the distance transcribed past the 3’ SS for nano-COP reads from control and U2 snRNA KD. Two biological replicates per condition are displayed. C) Global splicing order in nano-COP data from control and U2 snRNA KD for pairs of consecutive introns in reads that span two introns or more. Individual dots represent biological replicates. D) Distribution of total RNA splicing index for introns that are normally excised (“other”) or that are retained upon U2 snRNA KD, in the control and U2 snRNA knockdown conditions. Three biological replicates (“rep”) per condition are shown. The number of introns (N) in each category is indicated. E) Proportion of genes with and without intron retention in short-read RNA-seq from total RNA upon U2 snRNA KD, as a function of the exon skipping level in the same genes. The exon skipping inclusion level difference is defined as the difference in exon inclusion level between control and U2 snRNA KD. F) Distribution of the number of reads with SE, RI, both or neither in direct RNA sequencing data from control or U2 snRNA KD, for all the genes included in the SE and RI co-occurrence analysis shown in Figure 4D. G) Number of genes showing co-occurrence of SE(s) and RI(s) in transcriptome-wide cDNA-PCR nanopore sequencing data from WT and *NMF291^-/-^* (U2 snRNA mutant) mice^48^. The x-axis categories are the same as in Figure 4D. H) Example of a SE and RI co-occurrence in *Puf60* in *NMF291^-/-^*mice. The frequency of intron 6 being retained is higher given that exon 9 is also skipped (frequency_RI | SE_) compared to the total frequency of intron 6 being retained (frequency_RI_) (left plot), while the frequency of exon 9 being skipped is higher given that intron 6 is also retained (frequency_SE | RI_) compared to the total frequency of exon 9 being skipped (frequency_SE_) (right plot). ***: p-value < 0.001 in the one-sided binomial test assessing co-occurrence of SE and RI. In A) and D), boxplots elements are shown as follows: center line, median; box limits, upper and lower quartiles; whiskers, 1.5x interquartile range; points, outliers.

**Extended Data Figure 6 (related to Figure 4).**
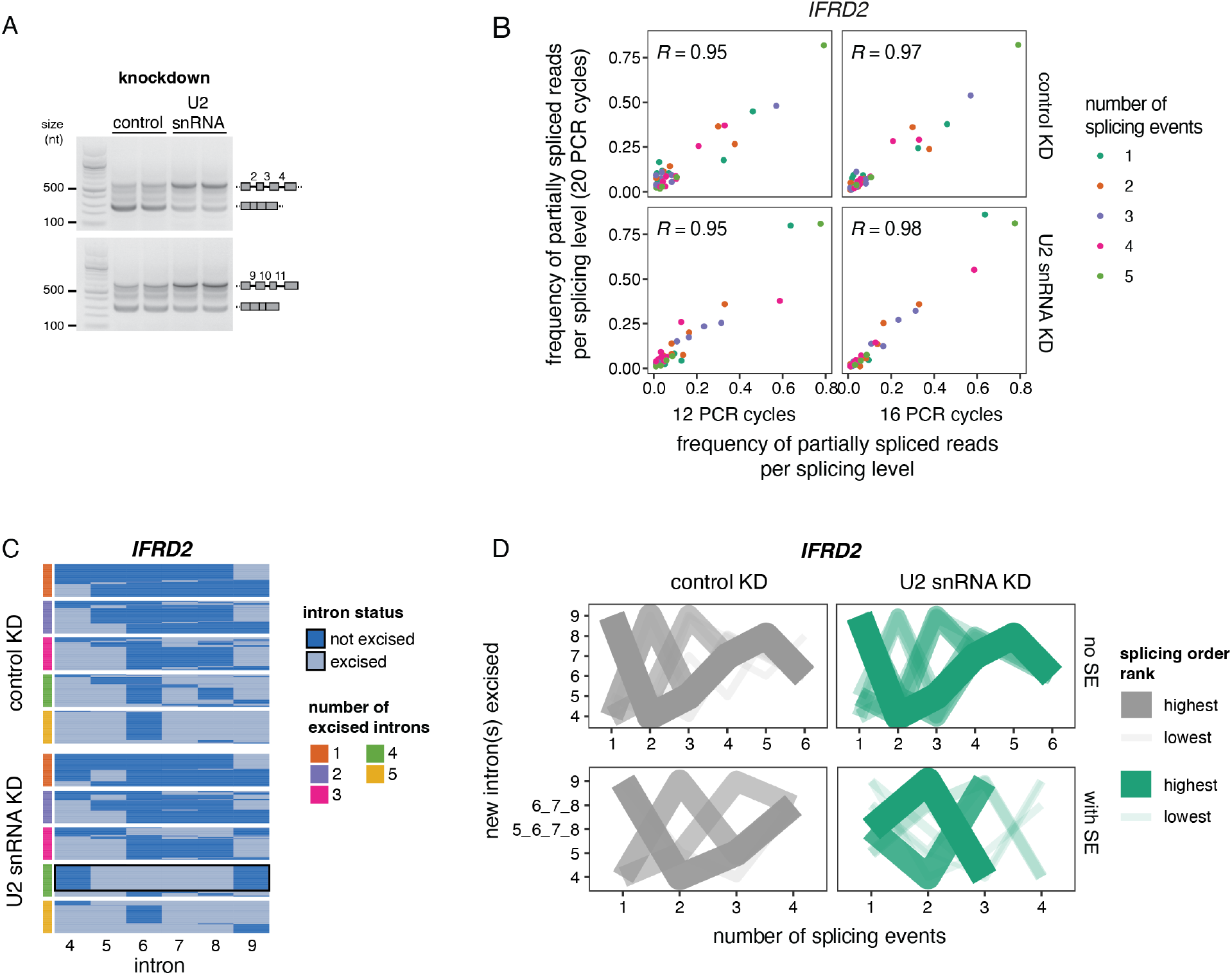
A) Agarose gel electrophoresis of the RT-PCR products from amplification of exons 2 to 5 and 9 to 12 of *IFRD2*, showing increased retention of introns 2 to 4 and 9 to 11 upon U2 snRNA KD compared to control. The identity of the spliced and unspliced products is shown on the right of the gel. Two biological replicates per knockdown condition are displayed. B) Scatter plots showing the correlation in frequency of intermediate isoform reads per splicing level for cDNA-PCR sequencing of *IFRD2* exons 4-10 from chromatin-associated RNA with 12, 16 or 20 PCR cycles. Pearson’s R correlation coefficients are shown on each plot (p-value < 0.0001, 95% confidence interval: control KD, 12 cycles: 0.91-0.97; control KD, 16 cycles: 0.95-0.99; U2 snRNA KD, 12 cycles: 0.91-0.98; U2 snRNA KD, 16 cycles: 0.97-0.99). The data obtained from 20 cycles was used for subsequent analyses. C) Representation of reads mapping to introns 4 to 9 of *IFRD2* upon cDNA-PCR sequencing of chromatin-associated RNA from control or U2 snRNA KD. Each line represents one read and each column represents one intron, with light blue showing excised introns while dark blue indicates present introns. The side color bar shows the number of excised introns in the intron group for each read. For each number of excised introns, 100 reads were subsampled from the total dataset. A black rectangle highlights the splicing order reversal upon U2 snRNA KD. D) Splicing order plots for *IFRD2* in the absence (top) or presence (bottom) of SEs. The thickness and opacity of the lines are proportional to the frequency at which each splicing order is used, with the top ranked order per intron group and sample set to the maximum thickness and opacity. On the y-axis, introns that are removed together to result in SE are shown separated by an underscore. The top 10 splicing orders per category (with or without SE) are shown. Experiments shown in B), C) and D) were performed on biological duplicates and data is displayed for one replicate per condition.

**Extended Data Figure 7 (related to Figure 5).**
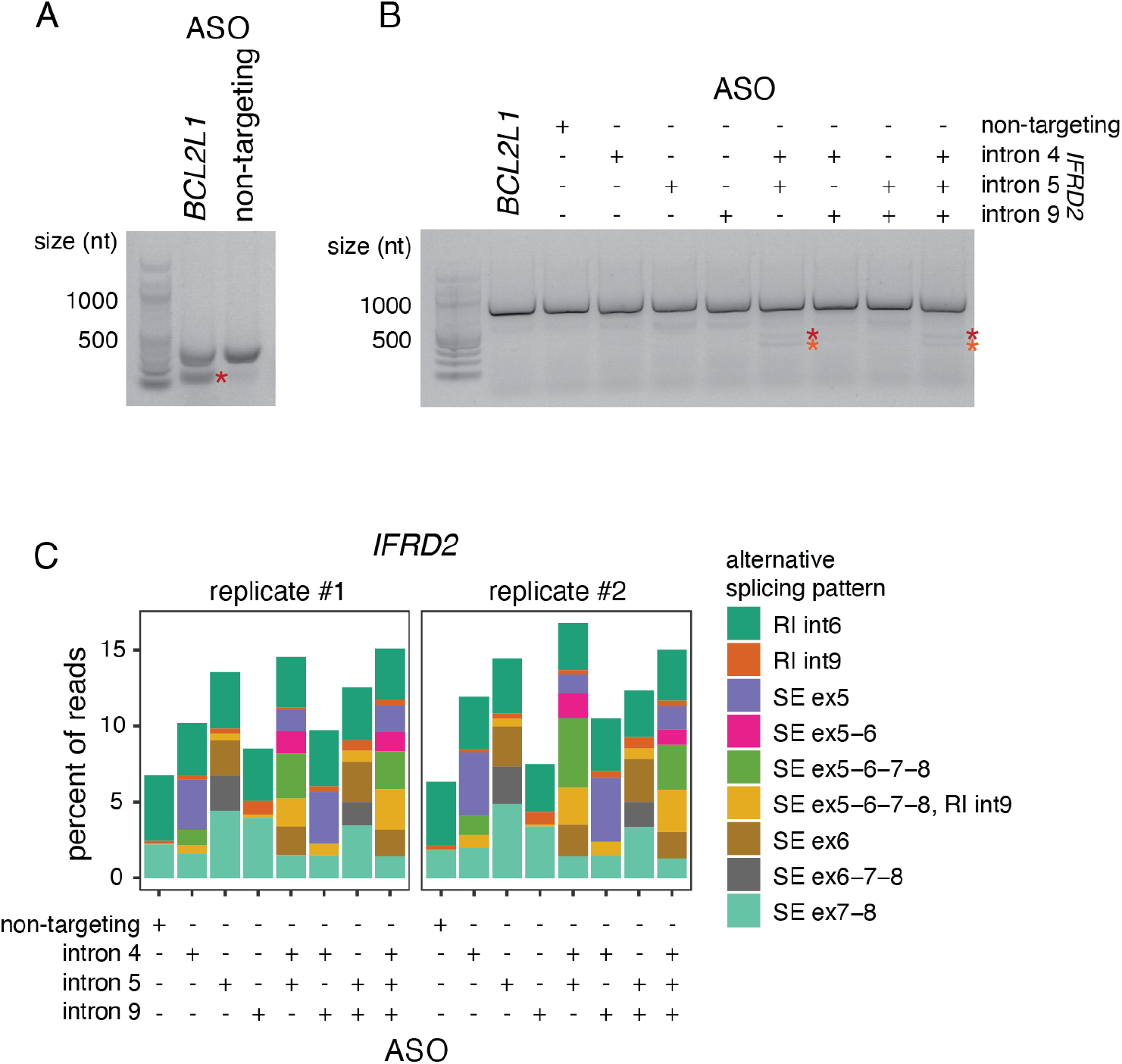
A) RT-PCR following 24 hour treatment with a positive control splice-switching ASO targeting a 5’SS in *BCL2L1*, compared to a non-targeting ASO. Treatment with the ASO results in the use of an alternative 5’SS that generates a smaller PCR product (red asterisk). B) RT-PCR following 24 hour treatment with ASOs targeting the 3’SS of introns 4, 5 and 9 of *IFRD2*. ASOs were used individually or in combination. Red and orange asterisks indicate the alternative splicing events in which exons 5 to 8 are skipped, without (orange) or with (red) intron 9 retention. C) Proportion of reads with different alternative splicing events observed in cDNA-PCR nanopore sequencing of total RNA following treatment with ASOs targeting *IFRD2*. RI: retained intron, SE: skipped exon(s), int: intron, ex: exon. Alternative intermediate isoforms present in at least 1% of reads are included. Intron 9 retention (RI_9) with or without skipping of exons 5-8 (SE_5-8) was included even when it was less frequent than this threshold.

**Extended Data Figure 8 (related to Figure 5).**
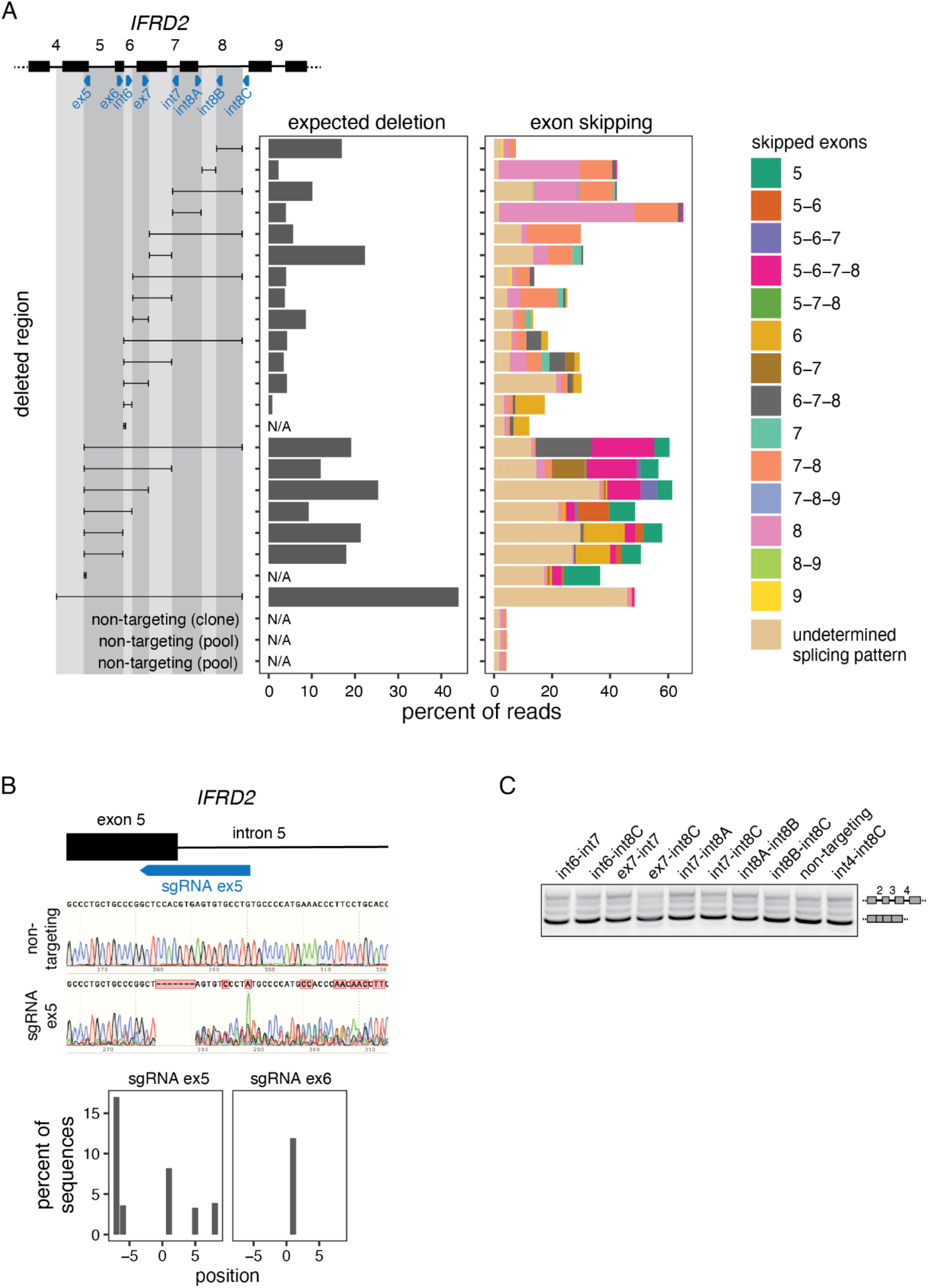
A) Deletion tiling of *IFRD2* using CRISPR-Cas9 editing and dual sgRNAs. Each sgRNA is represented as a blue triangle. For each individual sgRNA or sgRNA pair, the resulting deletion is shown as a black horizontal line delimited by vertical lines aligned to the sgRNA(s) used. Left: Percent of reads with the expected deletion in cDNA-PCR nanopore sequencing of chromatin-associated RNA. Deletions made with only one sgRNA and non-targeting controls were not assessed (N/A). Right: Percent of reads with exon skipping in each deletion. “Undetermined splicing pattern” refers to reads in which a splicing event does not map to annotated intron-exon junctions and results from detection of the deletion and/or from the use of a cryptic splice site as a result of the deletion. The reads shown on the left and right plots are not mutually exclusive, where reads containing the deletion (left) can also be classified as “undetermined splicing pattern” (right) if the deletion overlaps with exon(s). Pools of edited cells were used for this experiment, except for the largest deletion, which was found in a clonal cell line derived from cutting with the sgRNA int8-3 only. Non-targeting controls from pools of cells and a clonal cell line are also shown. Exon skipping events and undetermined splicing patterns present in at least 0.5% of reads are included. B) Top: Chromatograms from Sanger sequencing of pools of cells edited with a non-targeting sgRNA and one targeting the end of exon 5 as shown on the schematic. The bottom bar plot shows the type of indel introduced, as predicted from the chromatograms using TIDE^55^, for editing with sgRNAs against exon 5 (left) or exon 6 (right). The x-axis indicates the position relative to the expected cut site, which is 3 nt from the 3’-end of the sgRNA. C) Agarose gel electrophoresis showing the RT-PCR products from amplification of introns 2 to 4 of *IFRD2* in the deletions shown in D) that are not displayed in Fig. 5F. The identity of each RT-PCR product is shown on the right of the gel.

**Extended Data Figure 9 (related to Figure 5).**
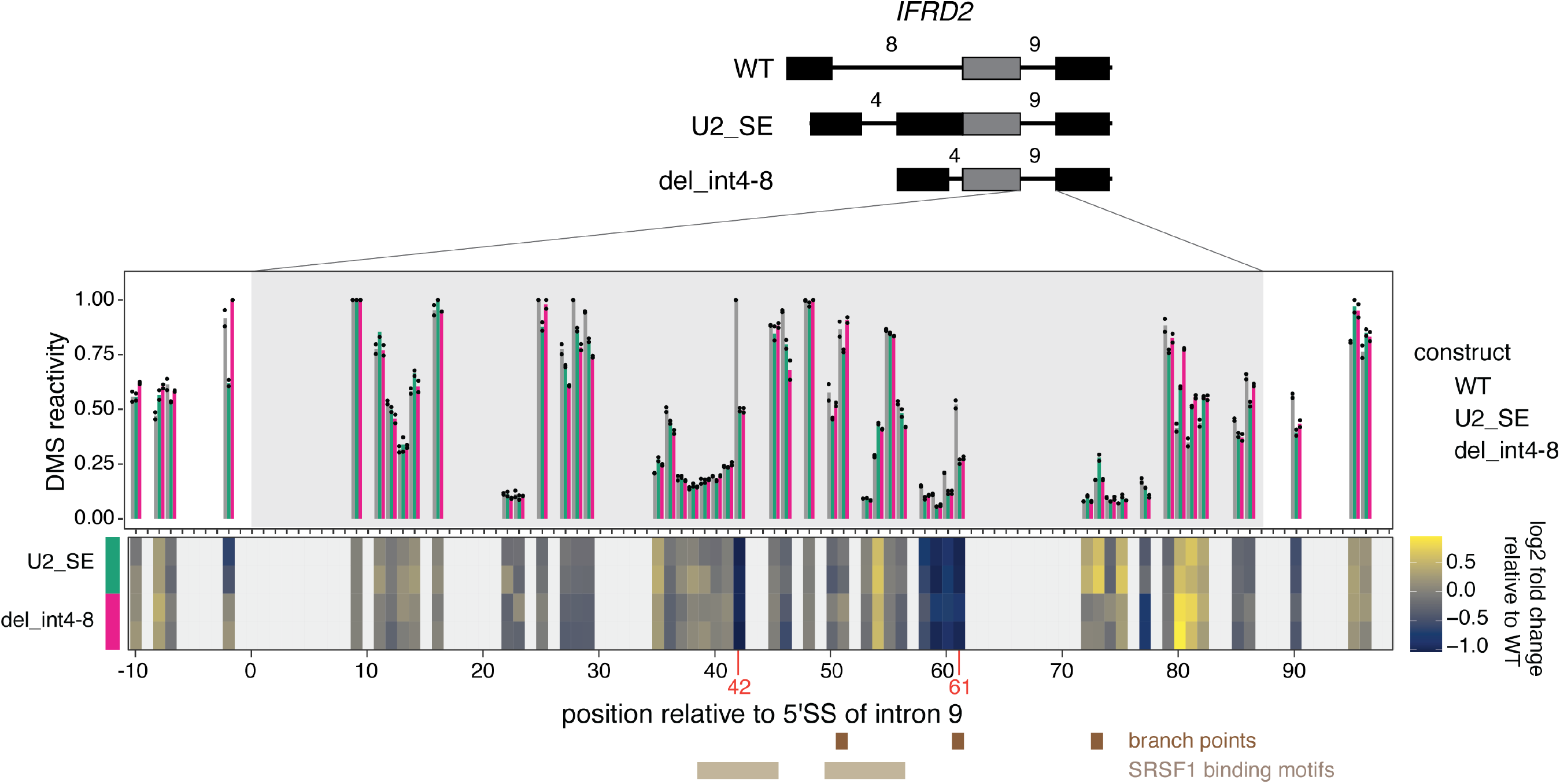
Top: DMS-MaPseq reactivity for *IFRD2* intron 9 +/- 10 nucleotides in the *in vitro* transcribed RNAs shown above. Individual dots represent two biological replicates. Bottom: Heatmap showing the log2 fold change of DMS reactivity in U2_SE or del_int4-8 relative to WT. Positions corresponding to G’s and T’s, which do not react with DMS, are shown in light grey. Positions where the largest fold change is observed in both mutated constructs are shown in orange font. Predicted branch points^41,54^ are shown as brown rectangles and the two predicted SRSF1 binding sites (ESEFinder^42^) with the highest scores are shown as beige rectangles.

## Acknowledgements

We thank members of the Churchman lab, W. Timp, D. Whye and D. Wood for helpful discussions, advice, and assistance; C. Patil, H. Merens, R.S. Isaac and N. Kramer for critical reading of the manuscript; D. Meng and Y. Jia for raw nanopore sequencing files from *NMF291*^-/-^ mice; the Biopolymers facility at Harvard Medical School and the Harvard University Bauer Core Facility for sequencing services. This work was supported by the NIH (R01-GM136794, R21-HG011682 and R01-HG010538 to L.S.C.), the Fonds de Recherche du Québec - Santé and the Canadian Institutes of Health Research (post-doctoral fellowship awards to K.C.). This research was conducted with support from the Human Neuron Core within the Rosamund Stone Zander Translational Neuroscience Center, Boston Children’s Hospital, which is also supported by the IDDRC (NIH P50HD105351).

## Author contributions

Conceptualization, K.C. and L.S.C.; Methodology, K.C. (lead), B.M.S., S.-L.D., S.R. and L.S.C.; Investigation, K.C. (lead), A.K., S.-L.D., B.M.S.; Software/Formal Analysis, K.C. (lead) and S.- L.D.; Writing – Original Draft, K.C. and L.S.C.; Writing – Review & Editing, K.C., A.K., S.- L.D., B.M.S., S.R. and L.S.C.; Funding Acquisition, K.C., S.R. and L.S.C.; Supervision, S.R. and L.S.C.

## Materials and Methods

### Cell culture

K562 cells (ATCC, CCL-243) were maintained at 37°C and 5% CO_2_ in RPMI 1640 medium (ThermoFisher, 11875119) containing 10% FBS (ThermoFisher, 10437036), 100 U/mL penicillin and 100 ug/mL streptomycin (ThermoFisher, 15140122). HeLa S3 cells (ATCC, CCL-2.2) were maintained at 37^°^C and 5% CO_2_ in DMEM medium (ThermoFisher, 11995073) containing 10% FBS, 100 U/mL penicillin and 100 ug/mL streptomycin.

### shRNA-mediated knockdown of EXOSC10

shRNA-expressing plasmids were obtained from Horizon Discovery (TRCN0000006338) or Addgene (scrambled control, #1864) (Supplemental Table 10). For lentiviral packaging, HEK293 cells were grown in six-well plates in antibiotic-free media and transfected with 500 ng shRNA plasmid, 500 ng psPAX2 (Addgene #12260), 50 ng pMD2.G (Addgene #12259) and 3.1 uL of FuGene HD (Promega, E2311) transfection reagent in 100 uL Opti-MEM medium (Fisher Scientific, 31-985-062). The media was replaced after 12-15 hours and viral media was collected 24 and 48 hours later. For spin transduction, 2 million K562 cells were resuspended in 1.75 media, mixed with 1.25 mL viral media and 8 ug/ml polybrene (Sigma TR-1003-G) and centrifuged at 1000g for 2 hours at 33°C. Cells were then pelleted by spinning at 100g for 5 minutes, resuspended in fresh media, and incubated at 37°C overnight. After 24 hours, 3 ug/ml puromycin (ThermoFisher, A1113803) was added to the media and selection was performed for 4 days, replacing the selection media every 48 hours. On day 4, cells were pelleted and resuspended in fresh media without puromycin. Cells were collected for cellular fractionation one day later. Total cell lysates to verify knockdown by qRT-PCR were also collected at the same time.

### Spinal motor neuron differentiation

Human induced pluripotent stem cells (iPSCs) were grown in 2D as a monolayer on a Vitronectin substrate in Stemflex culture medium until they reached a confluence of >90%. iPSC colonies were digested with Accutase for 5 minutes at room temperature until a single cell suspension was obtained and manual cell counts were performed using a hemocytometer. Suspensions were adjusted to a concentration of 1×10^6^ cells/mL and plated into Corning ultra low attachment dishes. On the day of dissociation (Day 0) cells were resuspended in a neural induction medium ‘N2B27’ with small molecules for dual Smad inhibition (SB431542-10uM and LDN 193189-0.25uM), WNT activation (CHIR99021-3uM), and a ROCK inhibitor (Y-27632-10uM) to generate adequate spheroid formation. The small molecule schedule continued as follows: Day 2-4 (SB431542-10uM, LDN193189-0.25uM, CHIR99021-3uM); Day 4-9 (Retinoic Acid-0.5uM, Smoothened Agonist-0.5uM, Purmorphamine-0.5uM); Day 9-14 (DAPT-10uM; Compound E-0.1uM; Culture One supplement-1x). Time points for collection occurred on days 0, 4, 9, and 14. Cells were dissociated using Accumax (Innovative Cell Technologies, AM105) for 15-20 minutes at room temperature. The reaction was inactivated with Ovomucoid (Papain dissociation kit, Worthington Biochemical Corporation). Cells were centrifuged for 5 minutes at 400g, resuspended in PBS, counted, and centrifuged again to pellet the cells before proceeding to cellular fractionation for purification of chromatin-associated RNA.

### Cellular fractionation

Cellular fractionation was performed as described in steps 6 to 19 of ^56^. For K562 cells, 8-10 million cells were collected and centrifuged for 5 minutes at 500g. At the end of the cellular fractionation, chromatin pellets were resuspended in 250 uL RIPA buffer (ThermoFisher, 89900) and mixed with 750 uL of Trizol LS lysis reagent (ThermoFisher, 10296028). For collection of nuclei, the protocol was stopped after step 14 and nuclei were resuspended in 250 uL RIPA buffer and mixed with 750 uL Trizol LS lysis reagent. For HeLa cells, up to 6 wells of a six-well plate (5 to 8 million cells) were washed with 1x PBS, scraped, collected, and centrifuged for 5 minutes at 500g. Cellular fractionation was performed as outlined above. At the end of the cellular fractionation, chromatin pellets were resuspended in 700 uL of Qiazol lysis reagent. For differentiating motor neurons, cellular fractionation was performed as outlined above with dissociated cells (6, 30-45 or 8-9 million cells/reaction for days 4, 9 and 14, respectively). At the end of the cellular fractionation, chromatin pellets were resuspended in 1 mL of Qiazol lysis reagent. Cell fractionations were verified by western blot as described in ^57^ with antibodies against RNA polymerase II CTD phospho Ser2 (Active Motif, 61984) and GAPDH (GeneTex GT239 or ThermoFisher MA5-15738). Cell fractionations from knockdown samples (EXOSC10 or U2 snRNA) were always performed in parallel with a scrambled control to account for small differences in the cellular fractionation that can occur with different batches of buffers and two biological replicates were collected for each condition.

### RNA extraction and reverse transcription

For total RNA extraction from K562 cells, cells were lysed in 250 uL RIPA buffer, incubated for 5 minutes on ice, and mixed with 750 uL Trizol LS. For total, chromatin-associated, and nuclear RNA extraction from K562 cells, 200 uL of chloroform was added to the samples in Trizol LS. Samples were briefly vortexed and centrifuged at 12,000g for 20 minutes at 4°C. The aqueous phase was transferred to a new tube, mixed with an equal volume of isopropanol and centrifuged at 20,000g for 20 minutes at 4°C. Pellets were washed with 950 uL of 75% ethanol, centrifuged at 20,000g for 5 minutes at 4°C, and resuspended in nuclease-free water. 2 ug of RNA was treated with DNase I (Invitrogen, 18068015). 700 ng of RNA was reverse transcribed using the SuperScript III First-Strand synthesis system (Invitrogen, 18080051) according to the manufacturer’s instructions. For total RNA extraction from HeLa cells and differentiating motor neurons, cells were resuspended in 700 uL Qiazol and incubated at room temperature for 5 minutes. Total RNA and chromatin-associated RNA from HeLa cells and total RNA from differentiating motor neurons were extracted using the miRNeasy kit (Qiagen, 217004) according to the manufacturer’s protocol, including the optional DNase treatment (Qiagen, 79254). 1 ug of RNA was reverse transcribed using the SuperScript III First-Strand synthesis system according to the manufacturer’s instructions. For chromatin-associated RNA purification from differentiating motor neurons, 200 uL of chloroform was added to the chromatin pellets in Qiazol. Samples were vortexed for 15 seconds and centrifuged at 12,000g for 15 minutes at 4°C. The aqueous phase was transferred to a new tube, mixed with an equal volume of isopropanol, incubated at room temperature for 10 minutes and centrifuged at 12,000g for 10 minutes at 4°C. Pellets were washed twice with 1 mL of 75% ethanol, centrifuged at 7,500g for 5 minutes at 4°C and resuspended in nuclease-free water.

### Poly(A) selection and nanopore direct RNA sequencing

Poly(A)+ RNA was purified using the Dynabeads mRNA purification kit (ThermoFisher, 61006) according to manufacturer’s instructions, starting with up to 40 ug of chromatin-associated or nuclear RNA. Direct RNA library preparation was performed using the kit SQK-RNA002 (Oxford Nanopore Technologies) with 500-700 ng of poly(A)+ RNA according to manufacturer’s instructions with the following exceptions: the RCS was omitted and replaced with 0.5 uL water and the ligation of the reverse transcription adapter was performed for 15 minutes. For most samples (see Supplemental Table 1), a yeast spike-in control was added to the libraries. Synthesis of yeast spike-in RNAs was based on the protocol described in https://www.ebi.ac.uk/ena/browser/view/PRJEB28423?show=reads for the *S. cerevisiae ENO2* gene. Six *S. cerevisiae* genes (*BDC1, ICT1, HIF1, ENO2, YKE4, HMS2*) were amplified from their genomic locus using HiFi Hotstart DNA polymerase (KAPA, KK2601) (primers in Supplemental Table 10) in a total volume of 100 uL using the following cycling conditions: 3 minutes at 95°C, then 30 cycles of 15 seconds at 95°C, 15 seconds at 62°C, 2 minutes at 72°C. The PCR amplicons were purified using 1X volume RNA Clean XP beads and eluted in 33 uL water. A second round of PCR was performed with nested primers, wherein the forward primer encodes a T7 RNA polymerase promoter site and the reverse primers have either 10, 15, 30, 60, 80, or 100 thymidines on the 5’ end (primers in Supplemental Table 10) using the following cycling conditions: 3 minutes at 95°C, then 18 cycles of 15 seconds at 95°C, 15 seconds at 62°C, 2 minutes at 72°C. The PCR amplicons were purified using 1X volume RNA Clean XP beads and eluted in 33 uL water. In vitro transcription was performed using 500 ng of DNA template and the MEGAScript™ T7 Transcription kit (Thermo, AM1334) according to the manufacturer’s instructions. RNA was cleaned up with the MEGAClear™ Transcription Clean-up kit (Thermo, AM1908) according to the manufacturer’s instructions, the concentration was measured by Nanodrop, and the size of the transcripts was verified by TapeStation. The six transcripts were pooled at an equimolar concentration (10 picomoles each). The spike-in was added at 5% of the amount of RNA used from the sample. Sequencing was performed for up to 72 hours with FLO-MIN106D flow cells on a MinION device in our own laboratory or with FLO-PRO002 on a PromethION device at the Harvard University Bauer Core Facility (Supplemental Table 1). Two biological replicates from K562 chromatin-associated RNA were sequenced. We also analyzed two of chromatin RNA replicates as well as two cytoplasmic and two total RNA samples (Supplemental Table 1) that were produced in ^58^. For all downstream analyses in K562 cells, replicates 3 and 4 produced in this study were combined into replicate A and replicates 1 and 2 produced in ^58^ were combined into replicate B (Supplemental Table 1) to increase coverage. For analysis of sMN differentiation, we sequenced biological duplicates for days 9 and 14, while we were limited to sequencing a single biological replicate for day 4 due to lower cell counts at this earlier timepoint. For total RNA from control and U2 snRNA knockdown, single biological replicates were sequenced on a PromethION device.

### RT-PCR and quantitative PCR

For RT-PCR, 1 uL of cDNA was used in a 20 uL reaction with Phusion polymerase (NEB, M03530). The following cycling conditions were used for PCR prior to agarose gel electrophoresis: 98°C for 1 minute, then 25 cycles of 98°C for 20 seconds, 60°C for 30 seconds, 72°C for 1.5 minutes, followed by a final extension at 72°C for 3 minutes. Electrophoresis was performed with a 1% agarose gel in 1X TAE buffer at 120V for 1 hour. For qRT-PCR, cDNA was diluted to 1:200 (snRNA KD) or 1:25 (motor neurons and EXOSC10 KD) and qRT-PCR was performed with Ssofast EvaGreen (Bio-Rad, 1725201) with the following cycling conditions: 95°C for 30 seconds, 40 cycles of 95°C for 5 seconds, 60°C for 10 seconds, and a final melt curve (65°C to 95°C with 0.5°C increment for 5 seconds). All primers used for PCR are listed in Supplemental Table 10.

### U2 snRNA knockdown

For U2 snRNA KD, 0.4 million HeLa cells were plated in six-well plates. After 24 hours, cells were transfected with 250 pmoles of antisense oligonucleotide (ASO)^22^ and 10 uL of Lipofectamine RNAiMAX (Fisher Scientific, 13-778-030) per well according to manufacturer’s instructions for forward transfection. The ASOs were ordered from Integrated DNA Technologies (IDT) and their sequences are indicated in Supplemental Table 10. Cells were collected 48 hours after transfection and resuspended in 700 uL of Qiazol lysis reagent (Qiagen, 79306) for total RNA extraction or carried forward to cellular fractionation for purification of chromatin-associated RNA.

### Illumina RNA-sequencing

mRNA-seq libraries were prepared with the mRNA HyperPrep kit (KAPA) in biological triplicates (control and U2 snRNA KD) and sequenced on an Illumina NovaSeq with paired-end 100 nt reads at the Harvard University Bauer Core Facility. Reads were trimmed using cutadapt v1.14^59^: cutadapt -a ${adapter_R1} -A ${adapter_R2} -m 20 -u 3 -q 20,0 --nextseq-trim=5 -- max-n 0 -o $out1 -p $out2 $fastq1 $fastq2. Reads were aligned to the human reference genome (build GRCh38) using STAR v2.5.4a ^60^ with parameters --outFilterMultimapNmax 101 -- outSJfilterOverhangMin 3 1 1 1 --outSJfilterDistToOtherSJmin 0 0 0 0 --alignIntronMin 11. Alternative splicing analysis was performed with rMATS-turbo^61^ (https://github.com/Xinglab/rmats-turbo). Exon skipping events were considered to be significantly more frequent upon U2 snRNA KD if they were supported by at least 10 reads in each replicate and had an FDR < 0.05. Exons with SE were binned by their inclusion level difference for the analysis shown in Extended Data Figure 5E. For intron retention, the splicing index was calculated using reads spanning each splice junction as: (number of spliced reads * 2) / number of unspliced reads, using custom scripts from ^14^ (https://github.com/churchmanlab/nano-COP). Splicing indices across conditions were compared using rmats-stat (https://github.com/Xinglab/rMATS-STAT) with a significance threshold of FDR < 0.05 and a splicing index fold change (U2 snRNA KD / control) > 2 or < -2. For calculation of splicing index in chromatin-associated short-read RNA-seq from K562 cells (Extended Data Fig. 2F), we used data from ^14^ (GEO accession GSE123191).

### NET-seq

NET-seq was performed as previously described in ^62^ in biological duplicates. Libraries were sequenced on an Illumina NextSeq with single-end 75 nt reads at the Harvard University Bauer Core Facility. Analysis was performed as described in ^62^ with the pipeline available at https://github.com/churchmanlab/MiMB2019NETseq.

### Nano-COP

Analysis of the distribution of 3’-ends in nano-COP data from K562 cells was performed with data from ^14^ (GEO accession GSE123191) as described in ^56^ with scripts available at https://github.com/churchmanlab/nano-COP. Nano-COP upon control or U2 snRNA KD was performed in biological duplicates as described in ^56^ but without 4sU labeling and selection. Briefly, cellular fractionation and extraction of chromatin-associated RNA were performed as described above following snRNA KD. Ribosomal rRNA depletion was carried out with Ribominus Eukaryote kit v2 (ThermoFisher, A15020). Poly(I) tailing and direct RNA nanopore sequencing were performed as described in ^56^ with 550 ng of RNA. Analysis of the distance transcribed past the 3’SS and of global splicing order for pairs of consecutive introns were performed as described in ^56^ with scripts available at https://github.com/churchmanlab/nano-COP.

### Nanopore sequencing of amplicons from cDNA

1 uL of cDNA was used in a 20 uL reaction with Phusion polymerase with primers targeting the region from exons 4 to 10 of *IFRD2* (Supplemental Table 10). The following cycling conditions were used: 98°C for 1 minute, then 12-20 cycles of 98°C for 20 seconds, 60°C for 30 seconds, 72°C for 3 minutes, followed by a final extension at 72°C for 3 minutes. 20 cycles were used for all experiments except for the comparison between 12, 16 and 20 cycles (Extended Data Fig. 6B). Two to three PCR reactions per sample were combined and cleaned-up using the Monarch PCR and DNA clean-up kit (NEB, T1030S). 10-100 ng of DNA from each sample was carried forward for end-repair using the NEBNext Ultra II End repair/dA-tailing Module (NEB, E7546S). The DNA was mixed with 1X volume of AMPure XP beads (Beckman Coulter, A63881), incubated on a rotator for 5 minutes at room temperature, washed twice with 70% ethanol and eluted in 22.5 uL water. For ligation of unique sample barcodes, each sample was combined with 2.5 uL native barcode (Oxford Nanopore Technologies, EXP-NBD104 or EXP-NBD114) and 25 uL of Blunt/TA Ligase Master Mix (NEB, M0367). The ligation reaction was incubated for 10 minutes at room temperature, mixed with 1X volume of AMPure XP beads, incubated on a rotator for 5 minutes at room temperature, washed twice with 70% ethanol and eluted in 12-26 uL water. After sample barcoding, an equal amount of DNA from each sample was mixed, for a total of 80-100 ng in 40 uL. This pool was combined with 5 uL of sequencing adapter AMII (Oxford Nanopore Technologies, EXP-NBD104 or EXP-NBD114), 50 uL of Quick Ligation Buffer and 5 uL of Quick T4 Ligase (NEB Quick Ligation Kit, M2200S). The ligation reaction was incubated for 10 minutes at room temperature, mixed with 0.5X volume of AMPure XP beads and incubated on a rotator for 5 minutes at room temperature. Sample clean-up and loading onto a FLO-MIN106D flow cell was performed with the Ligation Sequencing Kit (ONT, SQK-LSK109) according to manufacturer’s instructions. Sequencing was performed on a MinION device in our own laboratory.

### Splice switching with antisense oligonucleotides

For splice-switching ASO experiments, ASOs were ordered from IDT (Supplemental Table 10) with phosphothioate backbones and 2’O-methyl modifications on every base. We used an ASO targeting the 5’SS of *BCL2L1* as a positive control (Extended Data Fig. 7A)^63^ and an ASO targeting *HBB*, which is not expressed in HeLa cells, as a negative control^64^. 0.4 million HeLa cells were plated in six-well plates and transfected 24 hours later with 100 nM of ASO and 7.5 uL of Lipofectamine 3000 transfection reagent (Fisher Scientific, L3000008) per well according to manufacturer’s instructions for forward transfection. Cells were collected 24 hours after transfection and resuspended in 700 uL of Qiazol lysis reagent for total RNA extraction. All ASO experiments were performed on biological duplicates.

### CRISPR-Cas9 editing of IFRD2

sgRNAs targeting *IFRD2* were selected using CRISPOR^65^. Oligonucleotides corresponding to both strands of the sgRNA sequences were annealed and cloned into the plasmid lentiCRISPR v2 (Addgene #52961) as previously described^66^. For lentiviral packaging, HEK293 cells were grown in six-well plates and transfected with 2 ug lentiCRISPR v2 plasmid, 580 ng pRSV-REV (Addgene #12253), 1.16 ug pMDLg/pRRE (Addgene #12251), 700 ng pMD2.G (Addgene #12259) and 8 ul Lipofectamine 3000 with 8 ul P3000 reagent in 200 uL Opti-MEM medium. Cells were incubated at 37°C and viral media was collected after 48 hours. HeLa cells were transduced in six-well plates with 1.5 mL viral media and 1.5 mL normal media for single sgRNA transductions or with 1.5 mL viral media from each sgRNA for dual sgRNA transductions. In both cases, polybrene was added at a final concentration of 8 ug/ml. Cells were incubated overnight at 37°C and the media was replaced after 24 hours. Puromycin selection (1 ug/ml final concentration) was started after 72 hours and continued for 7 days, with replacement of the selection media every 48 hours. For generation of clonal cell lines, cells were diluted to 0.7 cells/mL, transferred to 96-well plates, and incubated until clones were detected. Genomic DNA was isolated with Dneasy Blood and Tissue kit (Qiagen, 69504). For verification of editing efficiency by Sanger sequencing, 1 ng of genomic DNA was amplified by PCR in a 20 uL reaction with Phusion polymerase and the following cycling conditions: 98°C for 1 minutes, 5 cycles of 98°C for 30 seconds, 68°C for 30 seconds, 72°C for 1.5 minutes, then 5 cycles of 98°C for 30 seconds, 65°C for 30 seconds, 72°C for 1.5 minutes, then 5 cycles of 98°C for 30 seconds, 60°C for 30 seconds, 72°C for 1.5 minutes, followed by a final extension at 72°C for 7 minutes. Sanger sequencing was performed at the Biopolymers facility at Harvard Medical School. Chromatograms were visualized with SnapGene (Insightful Science; available at snapgene.com). For editing with single sgRNAs, editing efficiency was estimated using TIDE (https://tide.nki.nl/)55. For each sgRNA combination, single biological replicates were analyzed.

### In vitro DMS-MaPseq

IFRD2_WT, IFRD2_U2_SE and IFRD2_del_int4-8 constructs (Fig. 5E) were ordered as G-blocks (IDT, see Supplemental Table 10) and amplified by PCR using a forward primer containing a T7 promoter sequence (Supplemental Table 10) with the Advantage HF 2 kit (Takara Bio 639124) and the following cycling conditions: 94°C for 1 minute, 27 cycles of 94°C for 30 seconds, 60°C for 30 seconds, 68°C for 2 minutes. The PCR amplicons were *in vitro* transcribed using the MEGAscript T7 Transcription Kit (ThermoFisher, AM1334). RNA was purified using the RNA Clean & Concentrator-5 Kit (Zymo Research, R1015). 1 ug of RNA was denatured at 95°C for 1 minute and subsequently refolded by incubating in 340mM sodium cacodylate buffer (Electron Microscopy Sciences) and 5mM MgCl_2_ for 20 minutes at 37°C in a thermomixer. After refolding, dimethyl sulfide (DMS) (Sigma, 471577) was added to a final concentration of 1% and incubated for 5 minutes at 37°C shaking at 800 rpm and the reaction quenched with 60µl β-mercaptoethanol (Sigma). DMS modified RNA was purified using RNA Clean and Concentrator-5 kit and eluted in 10 ul water. 500 ng of modified RNA was prepared for sequencing using the xGen™ Broad-range RNA Library Prep Kit (IDT, 10009865) with a modified reverse transcription step, using the TGIRT™-III enzyme (Ingex) and 5mM DTT. The RT reaction was incubated at 20°C for 10 minutes, 42°C for 10 minutes, 55°C for 60 minutes and the RNA was then degraded by adding NaOH to a final concentration of 200mM and incubating at 95°C for 3 minutes. After neutralizing the NaOH with 200mM of HCl, the library preparation proceeded as per the manufacturer’s protocol. Libraries were sequenced on an iSeq-100 with paired-end 151 nt reads. Raw reactivities were calculated using the DREEM-pipeline as previously described^67^. DMS-MaPseq was performed in biological duplicates for each condition.

### Nanopore data processing

Live basecalling of nanopore sequencing data was performed with MinKNOW (release 9.06.7 or later). For direct RNA sequencing data, all reads with a basecalling threshold > 7 were converted into DNA sequences by substituting U to T bases prior to alignment. Reads were aligned to the reference human genome (ENSEMBLE GRCh38 (release-86)) using minimap2^68^ with parameters -ax splice -uf -k14. For cDNA-PCR sequencing, all reads with a basecalling threshold > 8 were aligned to the reference human genome using minimap2 with parameters -ax splice. Uniquely mapped reads were extracted as outlined in ^56^. Coverage tracks in Figures 1 and 3 and Extended Data Figure 1 were produced with pyGenomeTracks^69^.

### Estimation of poly(A) tail lengths

Poly(A) tail lengths were estimated using nanopolish v0.13.3^30^ for direct sequencing of chromatin-associated RNA from K562 replicate B, which includes a yeast spike-in control. Raw signal fast5 files were indexed with nanopolish index and poly(A) tail lengths were calculated with nanopolish polya using default parameters. Reads with the quality control flag ‘‘PASS’’ and with estimated tail lengths greater than 0 were used in subsequent analyses. Normalization of poly(A) tails with the yeast spike-in control was performed using a median of ratios approach modeled after the size factor calculation for differential gene expression in DESeq (Love et al., 2014). Poly(A) tail lengths from reads mapping to the yeast spike-in sequences were extracted and the median tail length per spike-in was calculated in each sample, followed by the geometric mean of medians across samples. The ratio of the median poly(A) tail length per sample over the geometric mean was then computed. Finally, the size factor was defined as the median of ratios across the six spike-ins in each sample. Poly(A) tail lengths from endogenous genes were divided by this size factor for each read, yielding the normalized poly(A) tail length.

### Calculating read counts per gene in direct RNA sequencing

To obtain read counts per gene, aligned reads were mapped to annotated genes from RefSeq hg38 using bedtools v2.27.1^70^ intersect with options -s -F 0.5 -wo -a $RefSeq_bed_file -b $bam, requiring that at least half of the read maps to a given gene. The number of reads per gene was calculated in R. For measuring the abundance of sMN differentiation markers (Extended Data Fig. 3C), gene counts were normalized using the DESeq median of ratios approach^71^. The geometric mean of read counts per gene was calculated across samples. For each gene, the ratio of the read count per sample over the geometric mean was then computed. Finally, the size factor was defined as the median of ratios across all genes in each sample. For each sample, the read count for each gene was divided by the size factor to obtain the normalized abundance.

### Determination of the excision status of introns

To determine the excision status of introns in direct RNA nanopore reads, we used our previously published custom strategy described in ^56^. Briefly, RefSeq annotations of intron coordinates were extracted from the UCSC table browser for hg38 and reads that overlap introns on the same strand were extracted using bedtools intersect^70^. For each read/intron pair, features of read CIGAR strings were extracted for the portion of the alignment mapping to the intron and 50 nucleotides surrounding the 5′ and 3′ SSs. Introns were determined to be “excised” if a splicing event (CIGAR string “N”) starting and ending within 50 nt of the 5’ and 3’SS was present and the size of this splicing event was within 10% of the annotated intron size. Introns were determined to be “not excised / present” if there was no evidence of a splicing event and there was greater than 50% coverage in the 50 nt surrounding each splice site and at least 75% coverage within the region of intron it maps to. Introns where the read started within the intron and that met these criteria for the 3’SS were also considered “not excised / present”. Introns were determined to be “skipped” if a splicing event overlapped with greater than 50% of the 50 nt surrounding each splice site. Introns were determined to be “skipped_5SS” if a splicing event overlapped with greater than 50% of the 50 nt surrounding the 5’SS, started and ended within 50 nt of the 3’SS and the portion of the intron within the splicing event was within 10% of the annotated intron size. Introns were determined to be “skipped_3SS” if a splicing event overlapped with greater than 50% of the 50 nt surrounding the 3’SS, started and ended within 50 nt of the 5’SS and the portion of the intron within the splicing event was within 10% of the annotated intron size. Introns within aligned reads that did not meet these criteria were removed from subsequent analyses. For analysis of introns flanking alternative exons during motor neuron differentiation, the steps above were repeated for the GENCODE V38 annotations of intron coordinates, which were extracted from the UCSC table browser for hg38.

### Identification of introns with post-transcriptional removal

Reads spanning at least two introns were extracted. Reads from both biological replicates were combined. For plotting the number of present introns per read (Fig. 1C), only reads with a combination of excised and not excised introns were considered and the total number of spanned introns and present introns per read were computed. For the fraction of post-transcriptional excision per intron (Fig. 1E), all introns covered by a minimum of 20 reads were considered and the fraction unspliced was calculated as the ratio of unspliced reads by the sum of unspliced and spliced reads, after filtering out reads with undetermined or skipped statuses for the intron considered. For calculating the proportion of post-transcriptionally excised introns per transcript (Fig. 1F), transcripts covered by at least 20 reads were included. To determine the fraction of expressed transcripts and genes captured in our analysis, we used exonic RPKMs from short-read total RNA-seq data from K562 cells^14^ (GEO accession GSE123191). We defined transcripts/genes with RPKM >= 1 as expressed. The position (first, middle, last) of post-transcriptionally excised introns was determined from the intron annotation file (RefSeq hg38) and the intersection of intron positions per read was represented as an UpSet plot^72^ with the R package complex-upset (https://github.com/krassowski/complex-upset).

### Computation of splicing order

For direct RNA sequencing in K562 cells, groups of 3 or 4 introns from the RefSeq hg38 annotation were included in the analysis when they met the following criteria in both biological replicates: 1) each intron was present in at least 10 reads; 2) each splicing level, defined as the number of excised introns within each read for the considered intron group, was supported by at least 10 partially spliced reads that spanned all introns in the considered intron group. If a group of 3 introns was fully encompassed in a group of 4 introns, only the latter was kept for further analysis to avoid duplicates. For duplicated intron groups with the same genomic coordinates within different transcripts, only one instance was kept. For each splicing level *L*, the frequency *f_k_* of each possible intermediate isoform *k* was recorded by dividing the number of reads matching this intermediate isoform by the total number of reads at that splicing level. Next, we iterated through each level *L*, where for each observed intermediate isoform *k*, we identified the intermediate isoform(s) at the previous splicing level *L - 1* from which the isoform under consideration could originate (e.g. EXCISED_EXCISED_PRESENT could originate from PRESENT_EXCISED_PRESENT or EXCISED_PRESENT_PRESENT, see also Fig. 2B). Those intermediate isoforms were connected within a possible splicing order path and their frequencies *f_k_* were recorded. After iterating through each level, the frequencies of patterns supporting each possible splicing order *i* were multiplied to yield the raw splicing order score *P_i_*, where *N* is the total number of intermediate isoforms supporting a given splicing order (4 for groups of 3 introns and 5 for groups of 4 introns):

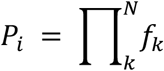

These raw scores *P_i_* were further divided by the sum of all raw scores for the considered intron group, where *n* is the total number of observed splicing orders for the intron group. This yielded the final splicing order score *p_i_* such that the sum of all scores *p_i_* was equal to 1:

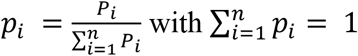

Only introns with “excised” or “not excised / present” statuses were considered for this analysis. For each intron group, Shannon diversity index (*H*) and evenness (*E*) were calculated as follows, where *p_i_* is the splicing order score for order *i*, *n* is the total number of observed splicing orders and *S* is the total number of possible splicing orders (6 for groups of 3 introns and 24 for groups of 4 introns):

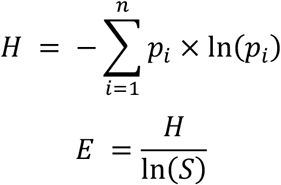

To identify reproducible splicing orders, we filtered for orders that showed the same rank in both biological replicates. For representation of splicing orders, we calculated normalized scores *np*, where the splicing order scores were divided by the maximum score for the intron group considered, such that the top ranked order had a score of 1:

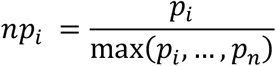

Splicing order plots were produced in R with ggplot2 (https://ggplot2.tidyverse.org) with the following command:

`ggplot(aes(x=splicing_level, y=new_intron_spliced)) + geom_line(aes(group=order_name, size=norm_score, alpha=norm_score))`

Splicing order plots in Figure 2, Figure 4 and Extended Data Figure 2 were made using data from one biological replicate A. Splicing order scores from both biological replicates are included in Supplemental Table 3.

For computing splicing order of constitutive introns during sMN differentiation, the same approach was used with the following modifications. Using intron annotations from RefSeq hg38, groups of 3 or 4 introns were included in the analysis when they met the following criteria in all samples: 1) each intron was present in at least 5 reads; 2) each splicing level was supported by at least 5 partially spliced reads that spanned all introns in the considered intron group. Only introns with “excised” or “not excised” statuses were considered for this analysis. For comparison of splicing order between different cell types and differentiation timepoints, intron groups present in all cell types being compared were included and the top splicing order for each cell type was extracted, with the following coverage thresholds: 1) each intron was present in at least 5 reads; 2) each splicing level was supported by at least 5 partially spliced reads that spanned all introns in the considered intron group. Intersections between each combination of cell types were represented using UpSetPlot (https://upsetplot.readthedocs.io/en/stable/).

### Simulation of random, SS-driven and “first-come, first-serve” splicing order

For each intron group used in the splicing order analysis in K562 cells, the number of reads mapping to each splicing level was extracted. At each splicing level, the corresponding number of introns were randomly removed without replacement. Splicing order was computed as described above using the randomly spliced reads. This was repeated 100 times and the average splicing order score per intron group across all iterations was recorded. For simulation of SS-driven splicing order, the same strategy was used but weights equal to 2^(5’SS_score+3’SS_score/)2^ were attributed to each intron during the random splicing step. 5’ and 3’SS scores were obtained from MaxEnt^53^. For simulation of “first-come, first-serve” splicing order, the same strategy was used but weights were attributed to each intron during the random splicing step based on the intron position from 5’ to 3’. For groups of 3 introns, the weights were 0.75, 0.23 and 0.02. For groups of 4 introns, the weights were 0.75, 0.15, 0.08 and 0.02.

### Identification of differentially abundant intermediate isoforms upon EXOSC10 KD

For each intron group analyzed in wild-type K562 cells, the number of reads matching each possible intermediate isoform was recorded. Intermediate isoforms with at least 10 reads in both biological replicates from at least one condition (scrambled or EXOSC10 KD) were kept for further analyses. A two-sided Fisher’s exact test was used to compare the frequency of the intermediate isoform considered and all other intermediate isoforms for the intron group between the two conditions. This was performed separately for each scrambled/EXOSC10 KD pair of biological replicates because of small differences in cell fractionation that can lead to more or less spliced reads between replicates. Multiple testing correction was performed using the Benjamini-Hochberg method. Intermediate isoforms were considered to be significantly upregulated in the EXOSC10 KD if FDR < 0.1 in both replicates and the odds ratio was > 1 in both replicates.

### Identification of AS introns during sMN differentiation

To allow for downstream analysis of splicing order, we used an intron-centric strategy to identify AS events during sMN differentiation. Intron annotations from GENCODE V38 were used since they contain a larger repertoire of alternative isoforms. Read intron statuses were combined from all samples and “not excised” introns were filtered out. For each intron, the proportion of reads with “SKP5SS”, “SKP3SS” or “SKP” status relative to the total number of reads was calculated. Introns with less than 50 total reads or less than 10 reads showing evidence of skipping were filtered out. Introns with a proportion of skipped reads between 0.05 and 0.95 were selected. From this list of introns, the proportion of skipped reads for each sample was calculated, and introns that had more than 10 skipped reads in at least one timepoint (in both biological replicates if applicable) were selected. A two-sided Fisher’s exact test was used to compare the frequency of skipped and spliced reads between two timepoints, for each biological replicate series. Multiple testing correction was performed using the Benjamini-Hochberg method. Introns were considered to be alternatively excised during sMN differentiation if at least two timepoints showed a difference of at least 10% in proportion of skipped reads and the comparison was statistically significant (FDR < 0.1) in both replicates. Of note, because many introns had high levels of “not excised” reads, in some cases only one intron flanking a cassette exon was considered to be alternatively removed, while in other cases both introns were considered to be alternatively excised. For duplicated introns with the same genomic coordinates within different transcripts, only one instance was kept for downstream analyses.

### Computation of splicing order with AS during sMN differentiation

For splicing order of AS introns, intron groups of 3 to 5 introns were defined from the AS introns identified in the previous section: If an intron alternative status was “SKP5SS”, the intron group was composed of this intron, the downstream intron and up to two upstream introns. If an intron alternative status was “SKP3SS”, the intron group was composed of this intron, the upstream intron and up to two downstream introns. If an intron alternative status was “SKP”, the intron group was composed of this intron and two flanking introns on each side. For duplicated intron groups with the same genomic coordinates, only one instance was kept. If a group of 3 or 4 introns was fully encompassed in a larger intron group, only the latter was kept for further analysis to avoid duplicates. Samples from all timepoints were combined and intron groups were included in the analysis if each splicing level was supported by at least 5 partially spliced reads that spanned all introns in the considered intron group. For computation of splicing order, we used the same strategy as described above, with the addition of one more type of splicing event: if the intron group included one “SKP5SS” and one “SKP3SS” introns that were consecutive or separated by introns with the status “SKP”, these introns were considered to be removed together as part of an SE event. For analysis of read patterns supporting AS splicing order at each timepoint (Extended Data Fig. 4D), intron groups were included if they were supported by at least two intermediate isoform reads that were consistent with the AS introns being removed last (AS introns present while other introns are excised) or being spliced earlier within the intron group (inclusion isoform: AS introns excised while other intron(s) are present, exclusion isoform: AS introns skipped while other intron(s) are present). The frequency of reads supporting AS introns being removed last was calculated by dividing the number of reads supporting AS introns being removed last by the sum of reads supporting AS introns being removed last or being removed earlier.

### Co-occurrence of exon skipping and intron retention

Intron excision status was determined as described above. To determine the inclusion status of exons, we used a similar approach: RefSeq annotations of exon coordinates were extracted from the UCSC table browser for hg38 and reads that overlap exons on the same strand were extracted using bedtools intersect^70^. Exons were determined to be “included” if the alignment file showed no indication of exclusion (CIGAR string “N” = 0) within 25 nt at the beginning and end of each exon, mapped portions of the read represent greater than 50% of the 25 nt at the beginning and end of the exon, and at least 75% of the read within the exon is aligned to the reference. Exons were determined to be “excluded” if there was no evidence of mapped sequence (CIGAR string “M” = 0) within 50 nt around both splice sites and the portion of the exon within the splicing event (CIGAR string “N”) was within 10% or 10 nt of the annotated exon size. If aligned reads that mapped to exons did not meet these criteria, the exon was classified as “undetermined” and excluded from subsequent analyses. For this co-occurrence analysis, introns were only considered if they were “excised” or “not excised”, while exons were only considered if they were “included” or “excluded”.

Short-read RNA-seq data was used to define a list of candidate genes with both SEs and RIs upon U2 snRNA KD: For SEs, all exons with at least 10 reads supporting the alternative event in each replicate, FDR < 0.05 and inclusion level difference > 0.01 were included. This low threshold was used since we noticed that some genes with low inclusion level difference with short-read RNA-seq had a higher inclusion level difference with nanopore direct RNA sequencing, likely due to some differences between the two technologies. For RIs, introns were included if they had FDR < 0.05 and splicing index fold change > 2. From these introns, those that were covered by at least 100 reads (excised or not) in each direct RNA sequencing sample were selected. Given the current length of nanopore sequencing, only introns < 1000 nt were considered. Genes that had at least one SE and one RI meeting these criteria were further analyzed. For each possible intron-exon pair within a read mapping to a candidate gene, intron splicing and exon inclusion statuses were recorded and the number of reads matching each possible status (“excised-included”, “excised-excluded”, “present-included”, “present-excluded”) were counted. For intron-exon pairs that showed at least 10 reads with the status “present-excluded” in the U2 snRNA KD sample, a two-sided Fisher’s exact test was employed to compare the frequency of this status to the status “excised-included” in control and U2 snRNA KD conditions. If the “present-excluded” pattern was significantly more frequent upon U2 snRNA KD (p-value < 0.05), the co-occurrence of the SE and the RI was assessed using two one-sided binomial tests. The first binomial test assessed whether the probability of the SE given the RI (SE | RI) was significantly higher than the frequency of the SE in all reads: scipy.stats.binom_test([counts(SE_RI), counts(RI)], p=freq(SE), alternative=greater). The second binomial test assessed if the probability of the RI given the SE (RI | SE) was significantly higher than the frequency of the RI in all reads: scipy.stats.binom_test([counts(SE_RI), counts(SE)], p=freq(RI), alternative=greater). Multiple testing correction was performed using the Benjamini-Hochberg method. Intron-exon pairs were classified into “both” if both binomial tests were statistically significant (FDR<0.05), “SE | RI” if only the first test was significant (FDR<0.05), “RI | SE” if only the second test was significant (FDR<0.05), or “not-significant” if neither tests were significant (FDR>=0.05). For analysis of splicing orders that contain introns involved in exon skipping and intron retention events, groups of three introns composed of the RI and the two introns neighboring the SE were defined. These intron groups were used to compute splicing order with chromatin-associated direct RNA sequencing data from K562 cells as described above, with one modification: only requiring that each splicing level be supported by 2 partially spliced reads that spanned all introns in the considered intron group in both replicates. To identify reproducible splicing orders, orders that showed the same rank in both biological replicates were included.

### Identification of AS and splicing order from cDNA-PCR nanopore sequencing

To determine the excision status of introns in cDNA-PCR sequencing reads, we used the same strategy as described above with RefSeq hg38 intron annotations. Since this data is not strand-specific, reads that overlap introns on either strand were extracted using bedtools intersect ^70^. Excision status was determined as above, with the exception that introns for which the read started within the intron were excluded. An intron-centric approach was used to identify AS events. Introns that were not excised were defined as “RI”. Introns were considered to be skipped if they were part of a series of consecutive introns with the most upstream having the status “SKP_3SS”, the most downstream having the status “SKP_5SS” and the middle introns having the status “SKP”. Exons downstream of the intron with status “SKP_3SS” and upstream of the intron with status “SKP_5SS” were considered to be “SE”.

For detection of expected CRISPR-Cas9 deletions from dual sgRNAs in cDNA-PCR sequencing reads, a similar strategy was employed. Deletions were defined starting from the expected cut site of the first sgRNA (3 nucleotides upstream of the PAM sequence) to the expected cut site of the second sgRNA. Deletions were determined to be present if a CIGAR string “N” started and ended within 50 nt of the 5’ and 3’ ends of the expected deletion and the size of this CIGAR string was within 10% of the expected deletion size.

For computation of splicing order, intron groups were defined as all introns included in the entire region amplified by PCR. The same strategy was used as described above, but without coverage thresholds. Splicing orders were computed once without SEs and once with SEs as follows: if the intron group included one “SKP5SS” and one “SKP3SS” introns that were consecutive or separated by introns with the status “SKP”, these introns were considered to be removed together as part of an SE event.

